# Characterizing drug activity with sensitive interactomes in human living cells

**DOI:** 10.1101/2025.11.20.689448

**Authors:** Cindy Kundlacz, Nawal Hajj Sleiman, Frédéric Marmigère, Benjamin Gillet, Sandrine Hughes, Laurent Gilquin, Hélène Duplus, Agnès Dumont, Gaël Yvert, Françoise Bleicher, Arnaud Gautier, Isabelle Coste, Toufic Renno, Samir Merabet

## Abstract

A number of human diseases results from abnormal protein-protein interactions (PPIs) involving key regulatory proteins. Therefore, an important strategy in therapeutics consists in developing inhibitory molecules that should ideally be specific for the aberrant PPI. In this context, it is critical to evaluate the number of PPIs that could be affected by the candidate molecule and to analyze the inhibitory potential before and after the formation of the PPI. Surprisingly, these two molecular aspects are rarely considered, due to a lack of appropriate methodological approaches.

In this study, we present a novel methodology that captures drug-sensitive PPIs by considering drug-induced cellular functions in live cell conditions. As a proof-of-concept, we identified interactions of the human core signaling protein ERK1 that are specifically affected by two different inhibitory molecules. In addition, we used a complementary set of innovative tools that allowed visualizing the inhibitory effect on ERK1/cofactor protein complexes after their assembly in living cells. Overall, our work establishes a unique methodological approach for deciphering drug activity for potentially any target bait protein of interest.

## Introduction

Cell-cell communication plays a pivotal role in determining cell fate and function and relies on diverse signaling molecules that bind to specific receptors in the plasma membrane. This receptor-ligand interaction initiates intracellular phosphorylation cascades, culminating in the activation of specific gene responses within the nucleus. One of the most widespread and conserved core phosphorylation cascade is the Ras-RAF-MEK-ERK (MAPK) signaling pathway, which is activated by receptor tyrosine kinase (RTK) receptors in response to various extracellular stimuli (e.g. growth factors, Fig. 1A) [1]. The Ras-RAF-MEK-ERK axis is involved in the regulation of fundamental cellular processes, including proliferation, growth, differentiation, migration and apoptosis. The central role of the MAPK signaling pathway is illustrated by the number of cancers (approximately one third) that result from its dysregulation in human [2]. Consequently, extensive research efforts have focused on identifying small molecule inhibitors targeting the Ras-RAF-MEK-ERK axis to counteract its aberrant activation in cancer [3,4]. The majority of these inhibitors primarily target RAF but their therapeutic application is limited by the observation of MAPK signaling reactivation and cancer resistance post-treatment [5,6]. Although fewer inhibitory molecules target the mitogen-activated protein kinase kinase family members MEK1 and MEK2 (thereafter denoted as MEK) or ERK1 and ERK2 (thereafter denoted as ERKs), there is growing consensus that combining RAF and MEK/ERK inhibitors could constitute a promising therapeutic strategy against the recurrence of MAPK-driven tumors [5,6].

**Figure 1.**
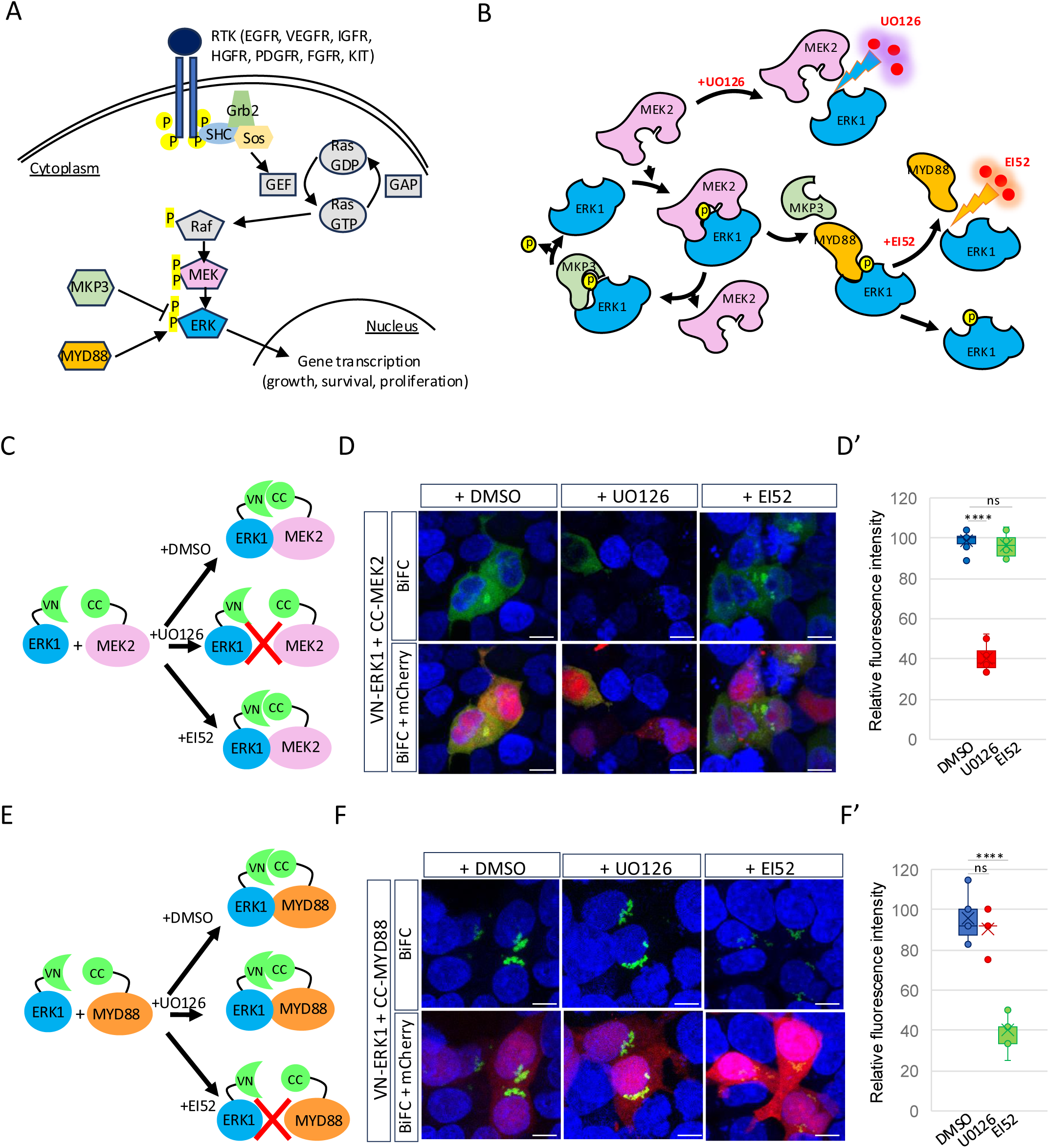
BiFC validation of the specific effect of U0126 and EI52 inhibitory molecules on their respective ERK1/Cofactor complexes in live HEK293T cells. **A.** Different ligands can activate the ERK pathway through the binding to a receptor tyrosine kinase (RTK). The phosphorylation of ERK by its upstream regulator MEK is inhibited by MKP3 and, reversely, protected by the binding to MyD88 [11]. **B.** The U0126 and EI52 inhibitory molecules affect the interaction between ERK1 and MEK2 or ERK1 and MyD88, respectively. **C-D’**. Validation of the effect of U0126 and EI52 on ERK1/MEK2 interaction, as assessed by BiFC in live HEK293T cells. **C.** Scheme of the different conditions. **D.** Illustrative confocal acquisitions of BiFC signals resulting from the interaction between VN-ERK1 and CC-MEK2 in live HEK293T cells, as indicated. **D’**. Quantification of the BiFC signal in the different conditions, as indicated. **E-F’**. Validation of the effect of U0126 and EI52 on ERK1/MyD88 interaction, as assessed by BiFC in live HEK293T cells. **E.** Scheme of the different conditions. **F.** Illustrative confocal acquisitions of BiFC signals resulting from the interaction between VN-ERK1 and CC-MyD88 in live HEK293T cells, as indicated. **F’**. Quantification of the BiFC signal in the different conditions, as indicated. All constructs were co-transfected with a mCherry-encoding plasmid (red) to normalize BiFC quantification with the transfection efficiency in each condition. Drugs were added in the transfection medium and present before the association between ERK1 and MEK2 or MyD88. Significance was determined relative to the mean BiFC value calculated with the control DMSO condition and was evaluated using t test (\*\*\*\**p < 0,0001*; ns, nonsignificant). VN: N-terminal fragment of Venus. CC: C-terminal fragment of Cerulean. Scale bar=10µm. See also Fig. S1 and Materials and Methods.

One of the first described MEK inhibitor is a chemical compound called U0126. It was identified from a screen for molecules that could suppress the expression of an AP-1 driven luciferase reporter in COS7 cells [7]. U0126 selectively blocks the activation of ERK by MEK (Fig.1B), thereby reducing cell proliferation in various cancer contexts [7–9]. Additionally, U0126 has also been reported to block interleukin production and reduce ERK-induced inflammatory response [10].

Another partner of ERKs, called MyD88, has been described to compete for binding with the phosphatase MKP3, allowing maintaining the phosphorylated state of active ERKs [11]. MyD88 was found to be overexpressed and to interact via its D domain with the D-recruitment site of ERKs in several cancer contexts, as well as to protect cancer cells against genotoxic agents [11]. Therefore, MyD88 represents a novel player in MAPK signaling for cell cycle control and cell transformation, constituting a promising target of choice for therapeutic development in MAPK-dependent cancers. Accordingly, an inhibitory molecule called EI52 has been developed to inhibit the interaction between the ERK1 isoform and MyD88 (Fig. 1B) and this molecule has recently been shown to trigger immunogenic apoptotic cancer cell death *in vitro* and elicit an anti-tumor T cell response *in vivo* [12].

U0126 and EI52 are initially described to affect two different PPIs (ERKs-MEKs or ERK1-MyD88), but their more complete molecular range of activity and specificity remains to be fully characterized. As a proof-of-concept, we proposed a novel experimental approach to analyze the activity of U0126 and EI52 on the ERK1 interaction potential with a specific gene library of approximately 900 different candidate human ORFs (Open Reading Frames). We called this library “ERKeome” as it contains genes involved in the regulation of the MAPK signaling pathway and genes of cellular functions induced by the two drugs, including apoptosis, immune pathway and inflammatory response. This work led to the identification of novel ERK1 interactions that are specifically inhibited by U0126 or EI52. Moreover, to better characterize the inhibitory potential of U0126 and EI52, we developed a complementary innovative method that allows visualizing their capacity in disrupting the target PPIs after their formation in living cells.

## Results

### Validation of the cell system

Our experimental strategy is based on the Bimolecular Fluorescence complementation (BiFC) technology, which allows visualizing protein-protein interactions in living cells. BiFC relies on the property of monomeric fluorescent proteins to be reconstituted from two separate hemi-fragments upon spatial proximity [13]. Given that our goal was to establish an ERKeome for performing a drug-sensitive screen, we first confirmed the suitability of BiFC in this perspective. More specifically, we tested whether BiFC could recapitulate the U0126- or EI52-sensitive interaction of ERK1 with MEK2 or MyD88, respectively. Practically, ERK1 was fused to the N-terminal fragment of the green fluorescent protein Venus (VN), while MEK2 and MyD88 were fused to the C-terminal fragment of the blue fluorescent protein Cerulean (CC, Fig. 1C and 1E). The VN and CC fragment can complement each other, producing a Venus-like fluorescent signal [14]. All constructs were placed under the same Doxycycline-inducible promoter (see Materials and Methods) and a red mCherry reporter vector was added for normalizing BiFC with transfection efficiency between the different conditions. Transfection of each construct in HEK293T cells showed clear distinct subcellular locations. VN-ERK1 was homogenously located in the cytoplasm and nucleus, underlining that it could exist as both inactive (non-phosphorylated in the cytoplasm) and active (phosphorylated in the nucleus) forms in HEK293T cells (Fig. S1). In contrast, CC-MEK2 was exclusively located in the cytoplasm, while CC-MyD88 concentrated in a few cytoplasmic foci juxtaposed to the nucleus (Fig. S1). These profiles were in accordance with previously described subcellular localizations or ERK1, MEK2 and MyD88 in other cell types [12]. Transfection of VN-ERK1 with either CC-MEK2 or MyD88 resulted in a BiFC fluorescent signal that reproduced the subcellular localization of MEK2 or MyD88, illustrating that the two cofactors were able to concentrate ERK1 at their respective specific subcellular locations (Fig. 1D and 1F). Importantly, pre-incubating the cells with U0126 before transfection (see Materials and Methods) led to a strong decrease of the ERK1/MEK2 BiFC signals, while no effect was observed for ERK1/MyD88 BiFC signals (Fig. 1D-D’ and 1F’). Inverse effects were observed with EI52, which affected ERK1-MyD88 but not ERK1-MEK2 BiFC signals (Fig. 1D-D’ and 1F-F’).

Altogether, these preliminary experiments confirmed that BiFC was appropriate to recapitulate the specific U0126- and EI52-sensitive interaction properties of ERK1 with the MEK2 and MyD88 cofactors in HEK293T cells.

### Designing a novel version of the Cell-PCA-based strategy with improved specificity and reproducibility

The original Cell-PCA (Protein Complementation Assay) strategy consisted in using a cell line expressing around 6000 human ORFs fused to the CC fragment for performing successive BiFC screens and comparing interactomes with different VN-fusion bait proteins in the same cellular context [15]. In the present work, we considerably modified the first version of Cell-PCA, and proposed a second generation (named Cell-PCAv2) that makes the approach more accessible, specific and reproductive.

The first parameter was the establishment of a biologically relevant CC-ORFeome, which contrasted with the previous large-scale design of Cell-PCA. Here, it consisted in generating a library of CC-ORFs covering functions related to cellular pathways that are regulated by ERK1 and modified upon the treatment with the U0126 or EI52 inhibitory molecules [10,12]. More specifically, we selected 984 genes associated with the MAP-K signaling and functions induced by U0126 and EI52 such as apoptosis, immune pathway and inflammatory response (Fig. 2A). These ORFs were used to establish a BiFC-ERKeome HEK293T cell line that eventually contained 907 integrated CC-ORFs after sequencing (Fig. 2A and Table S1, see also Material and Methods), of which 83 were described as ERK1 interactors in the BioGRID database (corresponding to 9,15% of the ERKeome; https://thebiogrid.org/111581/summary/homo-sapiens/mapk3.html and Fig. S2).

**Figure 2.**
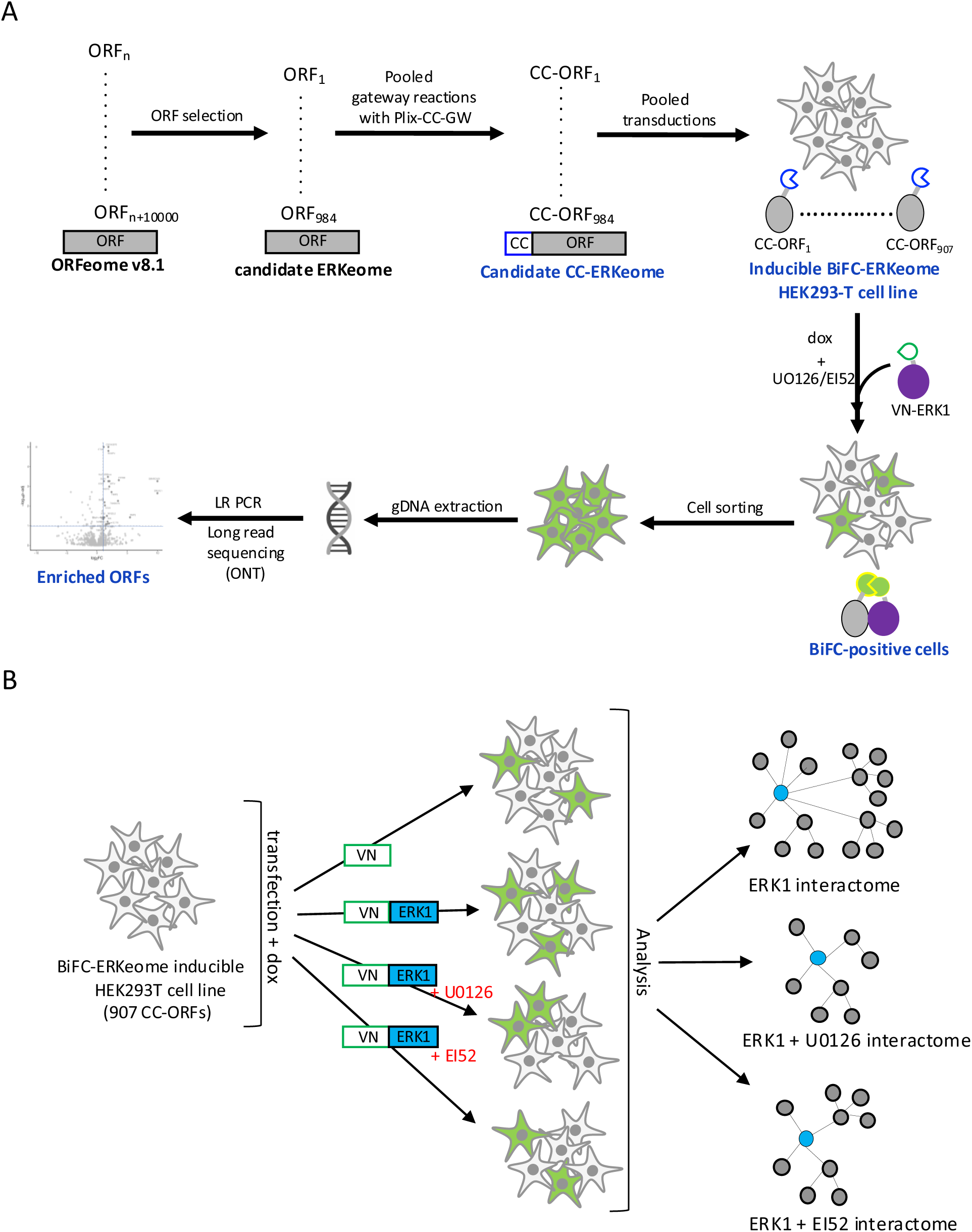
Principle of the Cell-PCAv2 strategy. **A.** Establishing a novel BiFC-ERKeome cell line to perform drug-sensitive BiFC screens with ERK1. Each step relies on new tools for improved specificity and robustness when compared to the original Cell-PCA approach [15]. See main text for details. **B.** Experimental conditions for identifying drug-sensitive interactions of VN-ERK1. VN: N-terminal fragment of Venus. See also Fig. S2-S3.

The second parameter that we modified was the expression system. A major drawback of Cell-PCA was the leakiness of the integrated CC-ORFeome (that was under the control of the Tet-off system, [15]), which prevented the maintenance of a stable cell line over several generations. Therefore, we built another vector where each CC-ORF is under the Tet-on promoter. This vector also expresses the most-recent generation of rtTA (rtTA-advanced) that is more sensitive to the doxycycline (dox) and has lower background than the original rtTA (Fig. S3). Altogether, these tools make the BiFC-ERKeome cell line highly stable over several generations, allowing performing successive screens with several biological replicates.

The third parameter implemented in Cell-PCAv2 was the selection procedure. In the first version of Cell-PCA, the selection of positive interacting ORFs was based on a frequency enrichment score when comparing the sorted fluorescent to the non-sorted cold cells. Here, BiFC screens were systematically performed with an additional condition corresponding to the VN fragment alone (Fig. 2B). This condition was considered as a reference threshold since the BiFC that could result from the association between VN and the CC-ORF gave a proxy of both the CC-ORF expression level and integration frequency in the BiFC cell line. Moreover, the comparison with VN alone allowed selecting interactions with a frequency score that was higher than the reference threshold, thus reflecting an increase in interaction affinity due to the specific interaction with the VN-bait fusion protein.

The last modified parameter was the identification of positive ORFs with a novel long-read sequencing protocol by Oxford Nanopore Technologies. It enabled getting the complete integrated CC-ORFs with less amplification cycles. The identification of positive CC-ORFs was achieved by establishing a novel analytical pipeline for automated application of statistics on read counts (see Materials and Methods).

### Identifying novel interactions of ERK1 with Cell-PCAv2 in live HEK293T cells

Before starting the cell sorting, we first validated that the BiFC-ERKeome cell line did not produce fluorescent cells either with dox, or upon transfection of VN-ERK1 in the absence of dox (Fig. S4). In contrast, the transfection of the VN fragment or VN-ERK1 construct induced a fluorescence in 5-6% of the whole cell population in the presence of dox (Fig. S4 and Materials and Methods). A total of 50 000 fluorescent cells was sorted in each condition and each experiment was performed in three biological replicates (see Materials and Methods). PCA analysis confirmed the correlation among the replicates and that the VN and VN-ERK1 conditions clustered separately (Fig. S5). We then applied a modified DESeq2 pipeline, which uses a model based on the negative binomial distribution allowing robust estimation of fold changes and significant differential enrichment, to automatically identify ORFs significantly enriched in association with VN-ERK1 when compared to VN alone. The selection was based on the adjusted p-value (equal to or less than 0,05 and calculated by considering all conditions: see materials and Methods) and on the log2FC (arbitrary set at a minimum of 0,75 to consider a significant enrichment when compared to the VN fragment). In total, 20 candidate cofactors met these selection criteria (Fig. 3A and Tables S2 and S3). Half of these candidate cofactors form a physical and functional cluster related to the MAPK signaling (Fig. 3B), which was also the most enriched GO term (Fig. 3C). Thus, although the BiFC-ERKeome was more enriched in GO terms related to the effects induced by U0126 and EI52, the interactome of ERK1 reproduced expected GO terms for the MAPK signaling in the absence of the two drugs. These observations underline that ERK1 preferentially interacted with specific cofactors of the candidate BiFC-ERKeome library. Finally, 5 out of the 20 positive candidates (25%) were already described as ERK1 interactors in BioGRID (Fig. S6), and the majority of the positive candidates (14/20) are predicted to contain an ERK-binding domain, thus reinforcing their potential to interact with ERK1 (https://scansite4.mit.edu/#home and Fig. S6).

**Figure 3.**
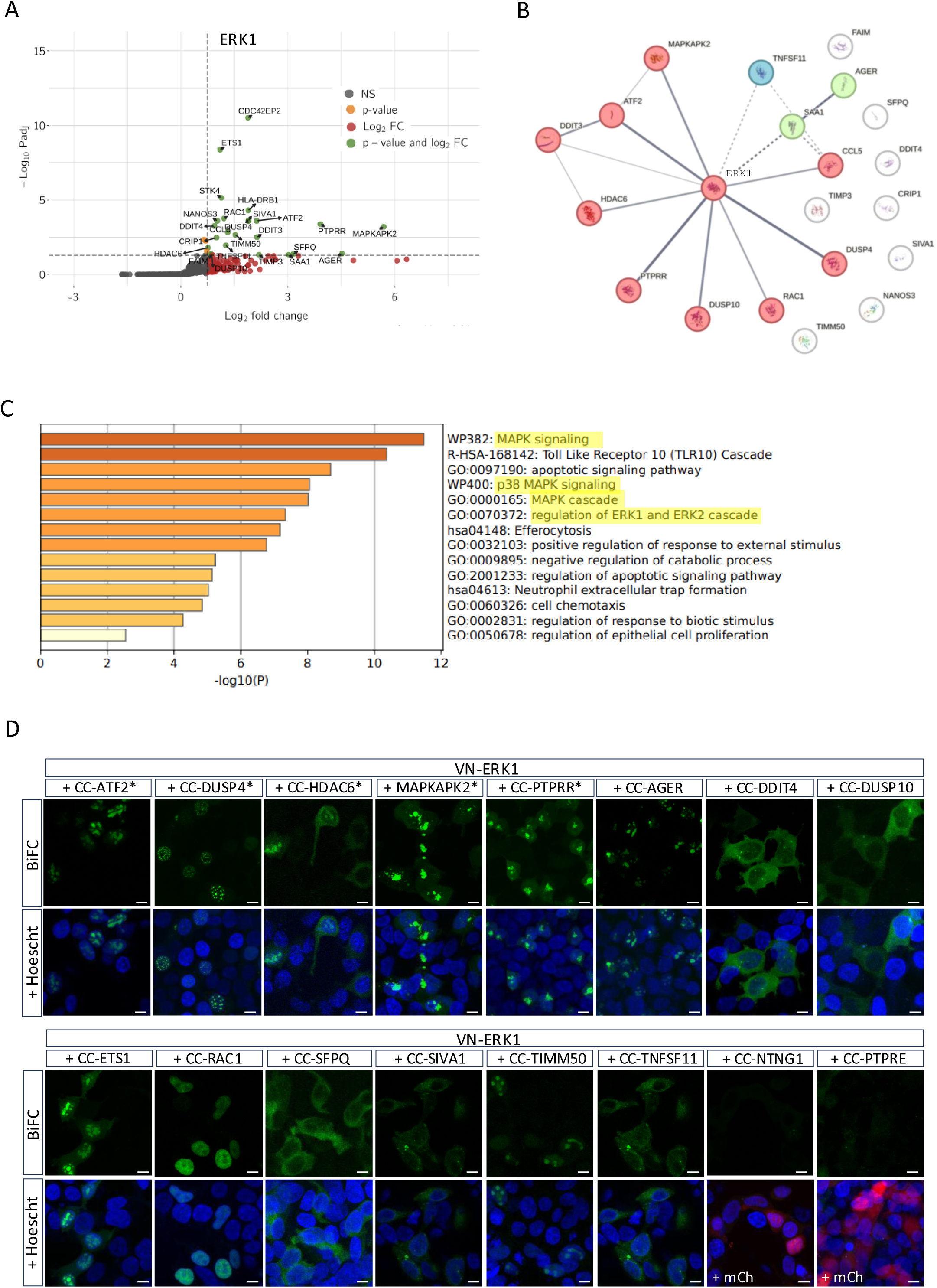
Capture of the ERK1 interactome with the Cell-PCAv2 strategy. **A.** Volcano plot of ORFs of the ERKeome distributed as a function of log2 fold changes (log2FC) and adjusted p-values when comparing the results between VN-ERK1 and VN. Selected interactions that met the log2FC (equal or superior to 0,75: dotted line on x axis) and adjusted p-value cut-off (in logarithmic scale, that is -Log10(0,05), dotted line y axis) are highlighted in green and labelled. Interactions that met only the adjusted p-value or Log2FC criterion are highlighted in orange or red, respectively. See also Materials and Methods. **B.** Clustering of ERK1-positive interactions with STRING. Interactions involved in the MAPK signaling pathways are highlighted in light red. Solid lines are proportional to the confidence score (the minimum required interaction score was set at medium confidence) and denote functional and physical interactions with ERK1 (see also Materials and Methods). **C.** GO-term enrichments as deduced from the ERK1 interactome (see also Materials and Methods). **D.** Individual BiFC assays confirming the ERK1-positive or ERK1-negative interaction status of selected ORFs from the BiFC screen. VN-ERK1 was individually transfected with the candidate CC-ORF and the interaction was analyzed in live HEK293T cells. Hoechst (blue) stains for the nucleus. The mCherry transfection reporter is shown only for the two BiFC-negative ORFs (NTNG1 and PTPRE). Pictures are illustrative confocal acquisitions from at least two different experiments. Scale bar=10µm. See also Materials and Methods and Fig. S4-S6.

To validate our list of potential ERK1-interacting proteins, we performed individual BiFC assays with 14 ERK1-positive (the 5 known ERK1 interactors and 9 novel interactions chosen to cover various functions) and 2 ERK-negative ORFs (randomly taken from the BiFC-ERKeome library) in living HEK293T cells (Fig. 3D). Interactions were all confirmed, and the analysis showed various and specific interaction profiles in the nucleus and/or cytoplasm, illustrating the diversity of contacts that could be engaged by the active or inactive form of ERK1 in HEK293T cells (Fig. 3D).

Overall, the screen with VN-ERK1 in the BiFC-ERKeome cell line revealed a limited number of positive interactions (20 out of 907), highlighting the specificity of the approach.

### Identification of novel ERK1 interactions sensitive to U0126 and EI52

We next analyzed the effect of U0126 and EI52 on ERK1 interaction properties in the same BiFC-ERKeome cell context. In parallel to the wild type condition, the screen with VN-ERK1 was performed by pre-incubating the BiFC-ERKeome cell line with U0126 or EI52 (Fig. 2B and Materials and Methods). The PCA analysis confirmed the distinct global effects of U0126 and EI52 on ERK1 interaction properties (Fig. S5). The same criterion of adjusted p-value (equal to or less than 0,05) and log2FC (set at a minimum of 0.75) was applied to select candidates that were significantly impacted by the drug treatment. In total, U0126 and EI52 affected 24 and 25 interactions, respectively (Tables S4 and S5). All these interactions were decreased when comparing their Log2FC with the wild type condition, highlighting that neither U0126 nor EI52 was able to induce ectopic interactions with ERK1. Among those interactions, we next considered more specifically the 20 ORFs found as positive interactors of ERK1 in the wild type condition. Under these filtering criteria, only three or two interactions could respectively be identified as U0126- (MAPKAPK2, SFPQ and TNFSF11) or EI52- (DDIT4 and TIMM50) sensitive (Fig. 4A-B). These results underlined that the two drugs had a relatively small range of action, with distinct effects on the ERK1 interactome.

**Figure 4.**
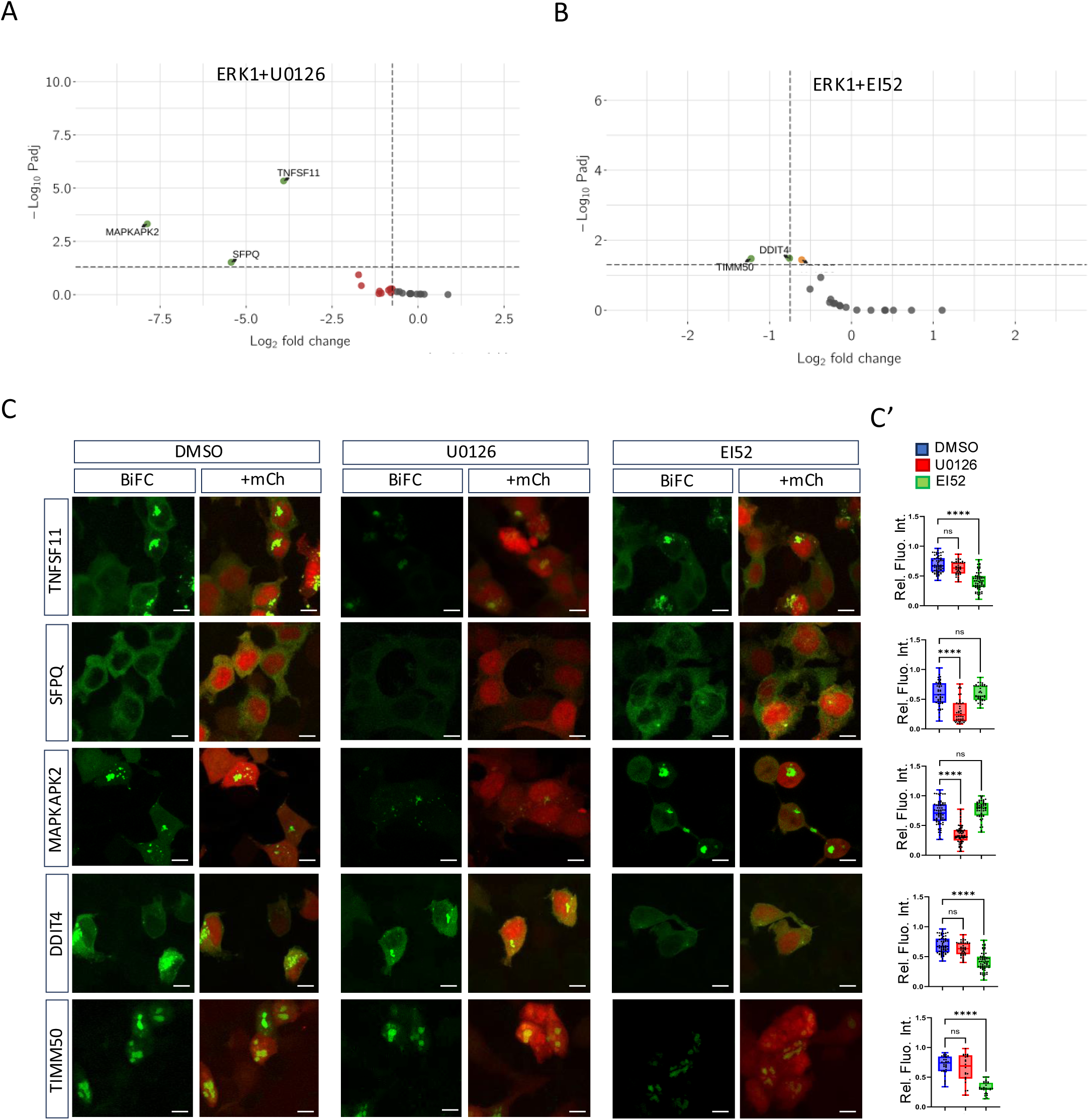
Effects of the U0126 and EI52 molecules on the ERK1 interactome. **A.** Volcano plot of U0126-affected interactions. **B.** Volcano plot of EI52-affected interactions. Significantly affected ERK1-positive interactions that met the log2FC (positive or negative, but equal or superior to 0,75) and adjusted p-value (equal or inferior to 0,05) criteria are highlighted in green and labelled. Interactions that met only the adjusted p-value or Log2FC criterion are highlighted in orange or red, respectively. **C.** Illustrative confocal acquisitions of individual BiFC assays between ERK1 and the selected drug-sensitive interactions. Constructs were co-transfected with the mCherry reporter for normalizing BiFC fluorescence between the three different conditions (DMSO, U0126 and EI52). Drugs were added before transfection. Scale bar=10µm. **C’.** Statistical quantification of BiFC signals (y axis: relative fluorescence intensity) in the different conditions, as indicated. Significance was evaluated using t test (\*\*\*\**p < 0,0001*; ns, nonsignificant).

Next, we decided to validate each drug-sensitive interaction by performing individual BiFC assays in the presence of one or the other drug. To this end, drugs were added at the same concentration as for the BiFC screen in the transfection medium. Quantifications confirmed the specific effect of UO126 and EI52 in each individual BiFC assay, thus validating the analysis from the BiFC-ERKeome (Fig. 4C-C’).

### Using the splitFAST2 system to analyze the inhibitory role of U0126 and EI52 after protein complex formation

Most of the current assays for validating the role of a candidate drug rely on treatments administered before protein complex formation. These assays provide information on the capacity of the drug to inhibit the initiation of the interaction, but not on its capacity to destabilize the protein complex once it is formed, which is an important parameter when considering the future therapeutic potential of a candidate molecule. This constraint applies particularly in the case of Cell-PCA, since BiFC is stabilizing the protein complex with the formation of irreversible interactions between the two non-fluorescent fragments. Therefore, BiFC is not appropriate for evaluating the capacity of a molecule to destabilize a dimeric protein complex after its formation.

To tackle this issue, we decided to switch to another complementation system called splitFAST2 that is reversible. The FAST (Fluorescence-Activating and absorption-Shifting Tag) protein can lead to instantaneous fluorescence upon non-covalent binding of fluorogenic 4-hydroxybenzylidene rhodamine (HBR) derivatives that are inherently non-fluorescent [16]. Several FAST variants have been engineered for using different chemogenetic reporters [17]. In addition, FAST can also be reconstituted upon the complementation of two sub-fragments, allowing analyzing PPIs in live cells [18,19]. The so-called splitFAST2 enzyme has recently been modified to display higher brightness and lower self-complementation [20].

The proof of concept and experimental parameters for using splitFAST2 were first established with ERK1/MEK2 and ERK1/MyD88 complexes, with or without U0126 and EI52 molecules (Fig. 5A and 5E). Here, the goal was to test the ability of U0126 and EI52 to destabilize their target protein complex post-formation. Practically, BiFC and splitFAST2 constructs were co-transfected in HEK293T cells and cultured for 12h without any drug, then incubated for additional 6h with the drug (or DMSO as a control) before assessing the effect on each protein complex. The HBR ligand was added just before live-cell imaging at the confocal microscope (see Materials and Methods). Control experiments with DMSO showed overlapping BiFC (green) and splitFAST2 (red) signals, confirming that these two different reporter systems did not affect interaction properties between ERK1 and either MEK2 or MyD88 (Fig. 5B and 5E, and Supplementary movies 1-6). BiFC and splitFAST2 signals were also not affected when incubating the cells with U0126 in the case of both ERK1/MEK2 (Fig. 5C and Supplementary movies 7-9) and ERK1/MyD88 (Fig. 5G and Supplementary movies 10-12) complexes. The result obtained with ERK1/MEK2 complex was not expected given that U0126 was shown to inhibit ERK1-MEK2 interaction when applied before dimeric complex formation (Fig. 1C’). Therefore, we concluded that U0126 could not destabilize ERK1/MEK2 complexes post-formation. Incubation with EI52 did not affect BiFC and splitFAST2 signals for ERK1/MEK2 complexes, as expected (Fig. 5D Supplementary movies 13-15). In contrast, EI52 led to an almost complete absence of splitFAST2 signals while BiFC remained unaffected in the case of ERK1/MyD88 complex (Fig. 5H and Supplementary movies 16-18). Thus, EI52 was able to destabilize the ERK1/Myd88 complex post-formation. Importantly, this result also confirmed that splitFAST2 was appropriate for revealing such effects.

**Figure 5.**
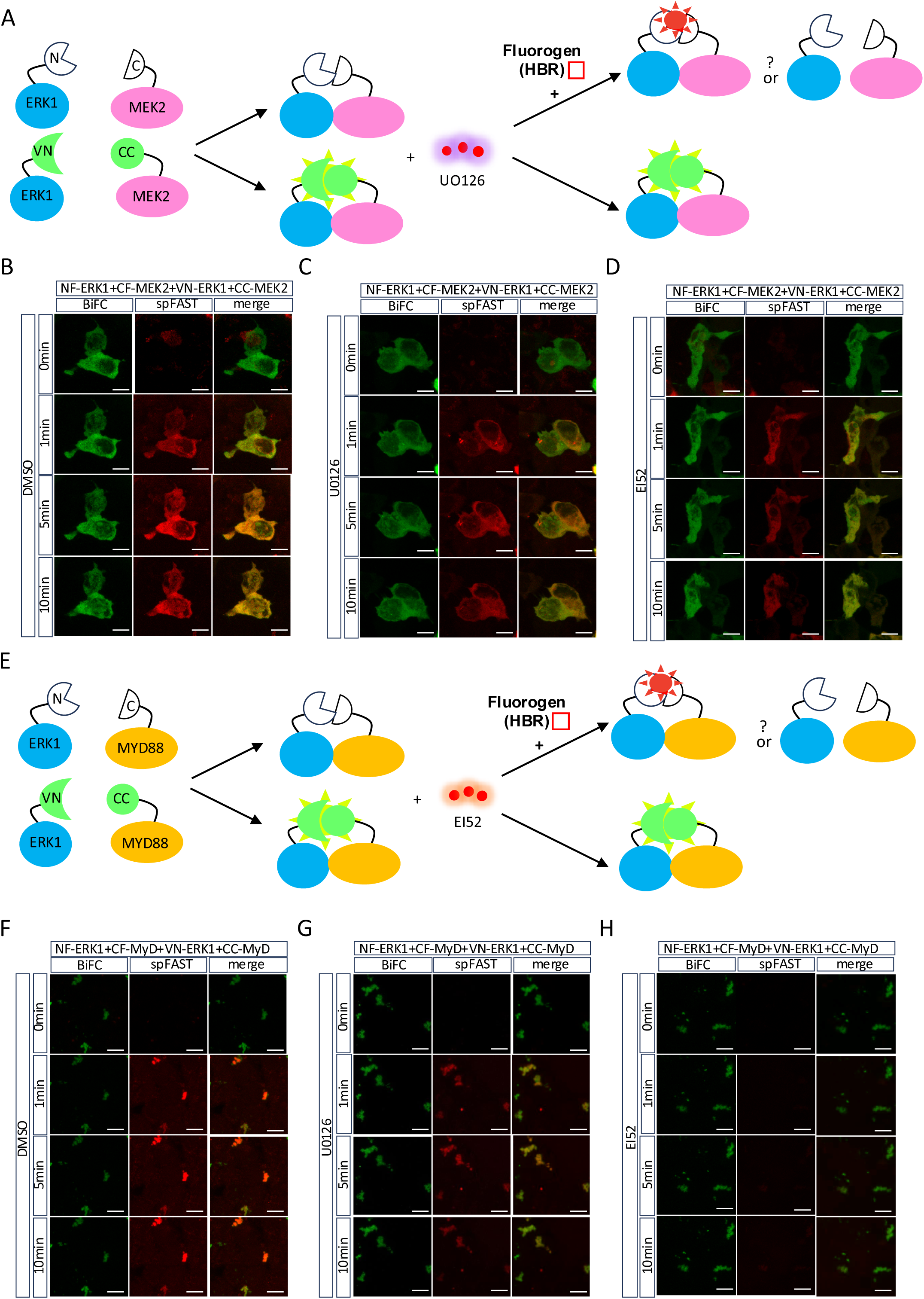
Establishing splitFAST2 for analyzing the effect of U0126 and EI52 on ERK1/Cofactor interactions post-complex formation. **A.** Scheme summarizing the potential effect of U0126 on ERK1/MEK2 interaction when assessed by doing BiFC and splitFAST2 and when tested after complex formation. The reversible nature of splitFAST2 allows assessing the potential destabilizing effect of U0126 post-complex formation. In contrast, the irreversible nature of BiFC does not allow assessing this level of inhibition. **B-D.** Illustrative confocal acquisitions of HEK293T cells transfected with the splitFAST2 (NFAST-ERK1 and CFAST-MEK2, red) and BiFC (VN-ERK1 and CC-MEK2, green) constructs before adding DMSO (B) or DMSO with U0126 (C) or EI52 (D), as indicated (see also Materials and Methods). Acquisitions are shown at 0, 1, 5 and 10 minutes (min) after the addition of the red HBR ligand. Neither BiFC nor splitFAST2 signals were affected in any condition. **E.** Scheme summarizing the potential effect of EI52 on ERK1/MyD88 interaction when assessed by doing BiFC and splitFAST2 and when tested after complex formation. The reversible nature of splitFAST2 allows assessing the potential destabilizing effect of EI52 post-complex formation. In contrast, the irreversible nature of BiFC does not allow assessing this level of inhibition. **F-H.** Illustrative confocal acquisitions of HEK293T cells transfected with the splitFAST2 (NFAST-ERK1 and CFAST-MyD88, red) and BiFC (VN-ERK1 and CC-MyD88, green) constructs before adding DMSO (F) or DMSO with U0126 (G) or EI52 (H), as indicated (see also Materials and Methods). Acquisitions are shown at 0, 1, 5 and 10 minutes (min) after the addition of the red HBR ligand. splitFAST2 signals were specifically affected upon the addition of EI52. Illustrative confocal acquisitions are from three independent experiments. Scale bar=10µm. See also Supplementary movies 1-6.

Next, we combined BiFC and splitFAST2 to analyze the effect of U0126 and EI52 in the context of the novel UO126- (ERK1-TNFS11, ERK1-SFPQ and ERK1-MAPKAPK2) and EI52-(ERK1-DDIT4 and ERK1-TIMM50) sensitive interactions found with Cell-PCAv2. As expected, BiFC was never affected in all cases (Fig. 6A-E). Surprisingly, none of the U0126-sensitive BiFC interactions were also affected when looking at splitFAST2 signals (Fig. 6A-C), showing that UO126 was not able to destabilize any tested PPI post-complex formation. In contrast, splitFAST2 signals resulting from ERK1/DDIT4 and ERK1/TIMM50 complexes were strongly affected in the presence of EI52 (Fig. 6D-E). Thus, unlike U0126, EI52 had the capacity to destabilize each tested interaction post-complex formation. This result highlights that protein complexes have different levels of sensitivity depending on the inhibitory molecule considered.

**Figure 6.**
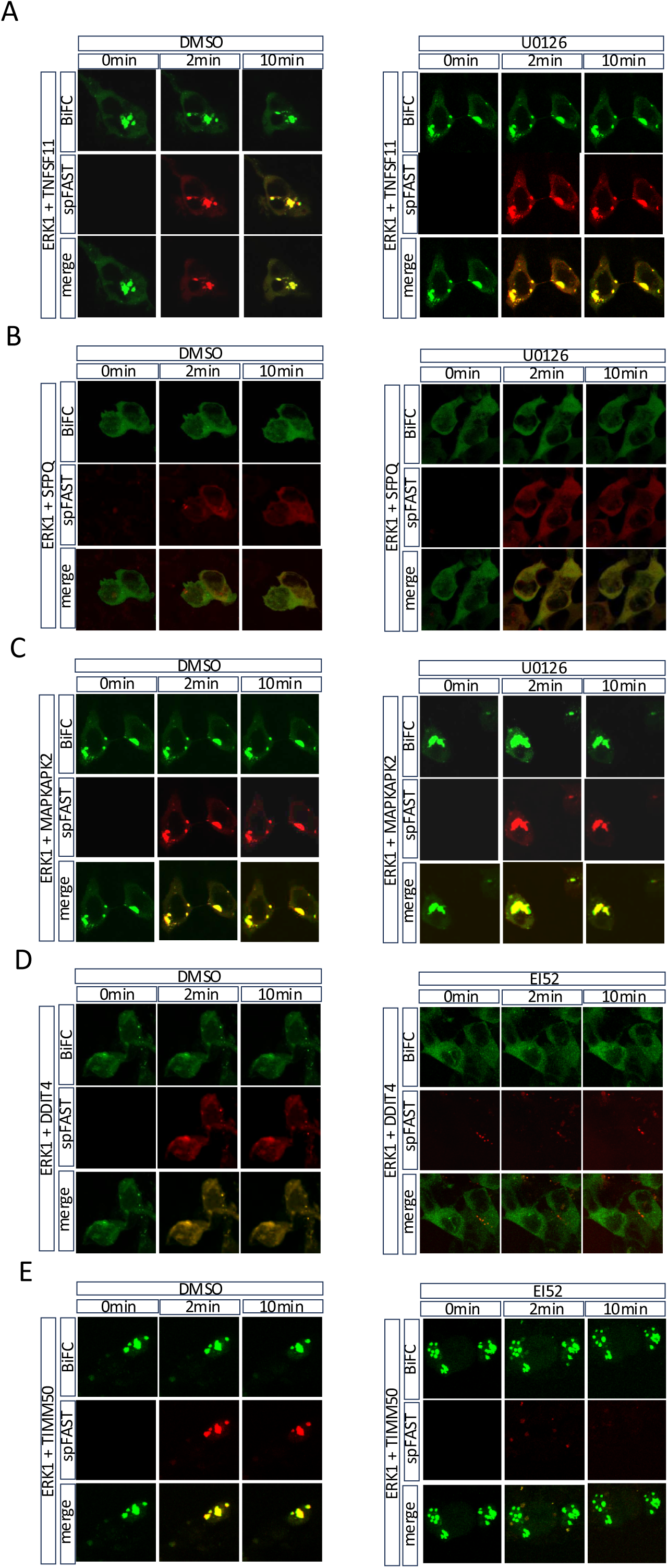
Testing the effect of U0126 and EI52 on the novel drug-sensitive interactions of ERK1 post-complex formation. **A.** Effect of DMSO or DMSO+U0126 on ERK1/TNFSF11 interaction. **B.** Effect of DMSO or DMSO+U0126 on ERK1/SFPQ interaction. **C.** Effect of DMSO or DMSO+U0126 on ERK1/MAPKAPK2 interaction. **D.** Effect of DMSO or DMSO+EI52 on ERK1/DDIT4 interaction. **E.** Effect of DMSO or DMSO+EI52 on ERK1/TIMM50 interaction. Cells were co-transfected with BiFC and splitFAST2 constructs and drugs were added as in Fig. 5. Illustrative confocal acquisitions are provided at the different times after the addition of HBR, as indicated. U0126-sensitive interactions were never destabilized post-complex formation while EI52-sensitive interactions were systematically destabilized. Each condition was performed in two independent experiments. Scale bar=10 µm.

## Discussion

### A novel Cell-PCA design for capturing drug-sensitive partners of ERK1 in human living cells

In this study, we presented an updated version of Cell-PCA [15] that relies on novel tools to make the approach more robust and specific. As a proof-of-concept, we established a BiFC cell line for screening new interactions of the ERK1 regulator in the context of drug-sensitive functions. To this end, the ERKeome was chosen to cover MAPK signaling pathway function and, importantly, to be enriched in physiological responses induced by the U0126 and EI52 molecules such as apoptosis, immune pathway and inflammatory response. The new screening strategy was based on a systematic comparison with the VN fragment alone and on several additional improvements, including a novel inducible expression system, a novel sequencing protocol and a novel statistical pipeline to select interactors with high confidence.

This work led to the capture of 20 interactions out of the 907 ORFs present in the BiFC-ERKeome cell line in the absence of U0126 and EI52. This low rate of positive interactions could be explained by several reasons. First, it reflects the specificity of the approach. Second, less than 10% of the 907 integrated ORFs have already been captured as ERK1 interactors by other approaches (https://thebiogrid.org/111581/summary/homo-sapiens/mapk3.html), highlighting that the drug-sensitive ERKeome did not cover many known ERK1 interactions. Third, the BiFC screen was performed under basal cell culture conditions for the MAPK signaling pathway, without inducing strong activation by adding growth factors for example. This parameter could explain that MyD88 was not captured although it was present in the BiFC-ERKeome cell line. Still, the most enriched GO-term among the 20 captured interactions was MAPK signaling pathway, although this GO-term was not the most enriched in the ERKeome.

Among the 20 ERK1-positive ORFs, half of them were described as direct ERK1 interactors and/or to functionally interact with the MAPK pathway in normal or oncogenic processes. In addition, the majority of positive ORFs (14/20) have a predicted ERK D-binding domain. Thus, our approach was successful in revealing novel ORFs as putative interaction partners of ERK1. These ORFs encode for proteins present in different cellular compartments, including the nucleus (such as AGER, ATF4, DUSP4, ETS1, HDAC6 or RAC1), the perinuclear region (PTPRR), the mitochondria (DDIT4 or TIMM50) or the cytoplasm (DDIT4, HDAC6 or SIVA1), which was also reflected in our individual BiFC assays with ERK-positive ORFs. Altogether, these observations underline that our cell expression system was sensitive enough to reveal diverse molecular and cellular functions of ERK1.

### Specificity of U0126 and EI52 inhibitory molecules

A key aspect in pharmacology is the molecular range of activity of cancer-inhibitory molecules for future therapeutic perspectives. In this study, we directly tackled this issue by constructing an ERKeome covering diverse cellular events induced upon treatment with U0126 and EI52. Surprisingly, very few interactions were affected by the two drugs and these effects were drug-specific. Among the interactions affected by U0126, only MAPKAPK2 was previously described as forming a complex with ERK1 [21,22]. SFPQ was recently proposed to regulate ERK activity during oocytes maturation [23] and TNFSF11 (also named RANK) was described to be activated by ERK during osteoclast differentiation [24]. None of these interactions has been correlated with cancer progression, which was not the case for the novel EI52-sensitive interactions captured in this work: DDIT4 has recently been shown to stimulate epithelial-mesenchymal transition in lung adenocarcinoma through the MAPK signaling pathway [25], and TIMM50 has been described to promote tumor progression via MAPK signaling in non-small cell lung cancers [26]. These observations underline that the two drugs are not equivalent in term of therapeutic potential. Importantly, the two molecules did not globally destabilize the ERK1 interactome, thus showing a certain level of specificity. Whether this level of specificity could apply to other ERK1-inhibitory molecules remains to be investigated. Here, we demonstrated that our experimental approach was appropriate for tackling this issue in the case of ERK1, and we propose that it could apply for any other bait protein that is the subject of pharmacological targeting development.

### Towards complementary complementation approaches for revealing inhibitory effects pre- and post-complex formation

A molecule with therapeutic potential is usually selected for its capacity to inhibit the formation of a specific PPI. Whether it could also destabilize the protein complex once it is formed is rarely considered. Still, this aspect is of critical importance when considering that (i) aberrant PPIs are by definition of stable nature and (ii) drugs are delivered post-complex formation *in vivo*. Therefore, testing this potential by using a reversible complementation system is necessary for selecting best potent molecules for future therapeutic perspectives. A unique reversible complementation system based on the split Nanoluciferase enzyme has been described in the literature [27], but this system has never been used for assessing complex disassembly upon drug treatment to our knowledge.

Here, we demonstrated that the splitFAST2 system was appropriate for revealing the destabilizing effect of EI52 on three different interactions (ERK1-MyD88, ERK1-DDIT4 and ERK1-TIMM50). In contrast, U0126 was inefficient to destabilize any of the PPIs that were sensitive prior to complex formation. These observations illustrate various levels of blocking activity therefore various levels of efficiency for EI52 and U0126. The finding that EI52 could destabilize ERK1-MyD88 interaction post-complex formation is in accordance with its previously described inhibitory role against MyD88-induced cancers [12]. It also illustrates that the affinity of EI52 is strong enough to replace previously established interactions with the D recruitment site of ERK. Moreover, the effect on ERK1-SFPQ and ERK1-TIMM50 complexes suggests that the anti-tumor activity of EI52 could also potentially rely on its blocking activity towards these two additional cancer-promoting interactions of ERK1.

In conclusion, we propose that combining our cell-based complementation methodology with splitFAST2 constitutes a straightforward approach for characterizing the molecular range of activity and specificity of candidate inhibitory molecules in the context of physiologically relevant interactomes for any target bait protein of interest.

## Materials and methods

### Cell lines

HEK293T and HEK293T-CC-ORFs cells were maintained in Dulbecco’s Modified Eagle’s Medium (DMEM-GlutaMAX-I, Gibco by Life Technologies) supplemented with 10% (v/v) heat-inactivated fetal bovine serum (FBS, Gibco), and 1% (v/v) penicillin-streptomycin (5000 U penicillin and 5 mg streptomycin/mL) at 37°C with 5% CO_2_. LentiX-293T cells were cultivated in Dulbecco’s Modified Eagle’s Medium (DMEM-GlutaMAX-I, Gibco by Life Technologies) supplemented with 10% FBS at 37°C with 5% CO_2_.

### Plasmids

A total of 984 ORFs from the V9.1 version of the hORFeome were fused at the 5’ end to the C-terminal part of the mCerulean gene (encoding the last 155-238 aa) using Gateway® technology. For individual BiFC assays, constructs were cloned into the plix _403 vector (a gift from David Root; Addgene plasmid # 41395; http://n2t.net/addgene:41395, accessed on 13 July 2020; RRID: Addgene_41395) and fused in 5’ end with either the N-terminal part of the Venus gene or the C-terminal part of the mCerulean gene. The VN-ERK and CC-ORFs constructs were placed under the same tet-off dox-inducible promoter. All constructs were sequenced verified before use.

### Establishment of the BiFC-ERKeome cell line

LentiX-293T cells were transfected with pLIX-CC-ORFs plasmids, the lentiviral packaging plasmid pCMVR8.74 and the lentiviral envelope plasmid pMD2.G using jetPRIME (Polyplus, Ref 114-15) and following the manufacturer’s recommendations. After harvesting the lentiviral particles, they were transduced in HEK293T at a low multiplicity of infection (MOI=0.3) to approximate one-gene-one-cell condition with ≥1,000X representation. During transduction, the culture medium was supplemented with 4 µg/mL polybrene (TR-1003, Sigma-Aldrich, Lyon, France) and replaced the next day. Two days later, selection of transduced cells was performed by adding 0.75 µg/mL of puromycin (Gibco, Cat No. A1113803) to the culture medium for one week, until the uninfected control cells completely died and the selected cells reached near confluency. The medium was changed daily during this selection process. The final amplified transduced cells were divided into aliquots of 1 x 10^6^ cells each and stored in liquid nitrogen for future screens.

### Cell-PCA screen

For each screen, a new batch of frozen BiFC-ERKeome cell line (1 x 10^6^ cells/ml) was cultured three days before the beginning of the experiment. Subsequently, BiFC-ERKeome cells were plated in 6-well plates (2.3 x 10^5^ cells per well). Two days later, at approximately 80% cell confluence, the cells were tranfected with either pLIX-VN-ERK (500 ng) or pLIX-VN-stop (500 ng) bait plasmids using 2 µL of jetPRIME buffer and 200 µL of jetPRIME reagent (Ref 114-15, Polyplus Transfection, France) per well. A mix containing jetPRIME buffer, jetPRIME reagent, plasmid DNA and 6 µM EI52 or 10 µM U0126 was incubated for 20 minutes at room temperature before transfection. During transfection, the medium was replaced with complete medium supplemented with 100 ng/mL dox. After 24 h, all transfected cells were pooled and resuspended in 200 µL PBS containing 2.5% SVF. 50,000 BiFC-positive cells were subsequently sorted using a BD FACS Aria II cell sorter (performed at AniRA-Cytometry core facility of the SFR Biosciences UAR3444/US8, Lyon, France). After each screening, the sorted cells were collected and genomic DNA was extracted using PureLink Genomic DNA Mini Kit (Cat No. K182001, Invitrogen, France) following manufacturer’s instructions. The extracted genomic DNA was then used for library preparation and was subjected to next-generation sequencing by in-house sequencing platform (PSI, IGFL, Lyon, France).

### Nanopore long reads sequencing and identification of the positive hORFs

DNA extracts were quantified using a Qubit 4.0. Primers were designed on plix plasmid sequence in order to specifically amplified CCORFs inserted in the pLIX plasmid. Forward primer was designed in the CC domain (5’ GACGAGCTGTACAAGAGATC 3’) and reverse primer in the hPGK promoter (5’ AACGGACGTGAAGAATGTG 3’) of the pLIX plasmid. CCORFs were amplified by Long Range PCR in 26 cycles using LongAmp Hot Start Taq 2X Master Mix (New England Biolabs). For each sample, the totality of the DNA extracted was amplified in a multiple tubes PCR approach (1 to 4 PCR tubes according to the quantity of DNA available per sample). In a final volume of 30µl, each PCR tube contained: 50ng of DNA, 15µl of Long Amp MasterMix and 1µl of each forward and reverse primer at 10µM. PCR conditions were as follows: initial denaturation step at 95°C for 1min, 26 cycles with: denaturation step at 95°C for 20 seconds, annealing step at 55°C 30 seconds and elongation step at 65°C for 4 minutes, final elongation for 5 minutes at 65°C. For each sample, all PCR tubes were pooled and purified using SPRI beads (Beckman Coulter) in ratio 1.7X. Final elution was performed in 30µl using ultra-pure water. Purified amplicons were quantified with Qubit 4.0 and quality assessed by TapeStation 4150 analyses (D5000 reagents and screentape; Agilent Technology). Oxford Nanopore Technologies (ONT) barcoded libraries were constructed according to the ONT Native Barcoding Amplicon protocol with the SQK-LSK109 kit and EXP-NBD104. To initiate the library construction, 200ng of amplified DNA per sample were used for End-Prep step. After native barcode ligation, maximum 24 quantified libraries (40ng per sample) were pooled in an equimolar manner. The pooled library was finalised by ligation of sequencing adaptor before being deposited on a flowcell R9.4.1 (FLO-MIN106) for sequencing with Mk1C device. Fastq reads were generated by MinKNOW 21.10.8 using guppy version 5.0.17 for basecalling. To analyze the data, ONT reads were filtered on their quality to retain the best quality reads (>Q12). A first BLAST approach against the pLIX plasmid was used to keep only those reads with the expected plasmid sequences at both ends. This step eliminates any artefactual reads resulting from PCR amplification of genomic fragments. A second BLAST approach was then performed using the hORFeome v8 reference database. Reads with a match on a CCORF of the database with at least 85% identity, and with an alignment length between 90 and 110% of the CCORF length, were assigned to the first BLAST hit (max_target_seq parameter set to 1). In this way, a read is assigned to only one CCORF. The BLAST parameters were chosen in order to take into account the amplification and ONT sequencing error rates. The BLAST results for all reads were then aggregated to generate a count table of CCORFs/genes detected per sample. In some cases, the parameters chosen for the BLAST approach are not able to differentiate between certain CCORFs. In order to identify these closest CCORFs, a BLAST of the hORFeome V8 DB against itself was performed using the same approach and parameters as for reads. All raw data are provided in Supplementary Table 6.

### Identification of ERK1-interacting ORF candidates (DESeq2)

All analyses were performed with DESeq2 (v.1.46.0), a package for differential expression analysis from the R programming language (v.4.4.0), included as part of the Bioconductor project (http://www.bioconductor.org/packages/release/bioc/html/DESeq2.html and [28]). DESeq2 utilizes a custom class object called DESeqDataSet, which stores the data required to perform the differential expression analysis. Two input files are required: the first containing the un-normalized counts data and the second containing a table with samples information. Clustering analysis was conducted to assess similarities between samples. A heatmap of the sample-to-sample distance matrix was created using the pheatmap and RColorBrewer packages. The darker the color, the closer the samples are to each other. The distance matrix computed on the transformed counts data, using the variance stabilizing transformation (VST) approach which aims to remove the dependence of the variance on the mean.

Complementary to the clustering analysis, a principal components analysis (PCA) [29] was applied to the transformed counts data to visually inspect the presence of batch effects. PCA is a dimension reduction technique that projects a dataset onto a subspace of lower dimension described by an orthogonal basis. The plotPCA function of the DESeq2 package can be used to project the VST data onto the 2D-plane formed by the first two principal components, namely PC1 and PC2. The percentage on each axis label represents the fraction of total variance explained by each principal component.

After obtaining the list of candidates from the DESeq2 test for differential expression, two filtering criteria were further applied to discriminate ORFs: a log2 fold change greater than 0,75 (log2FC ≥ 0,75) and an adjusted p-value smaller than 0.05 (P-adjusted ≤ 0.05).

A volcano plot was used to visualize the differentially expressed ORF candidates. Here, a custom function (lscript VolcanoPlot.R), based on the EnhancedVolcano package, was used to adapts the graph. The vertical line marks the log2 fold-change cut-off. The vertical line marks the adjusted p-value cut-off in logarithmic scale, that is -log10(0.05).

To reproduce the results of the DESeq2 analysis, an R script (main.R) is available on the following GitLab repository: https://gitbio.ens-lyon.fr/igfl/merabet/BIFCScreen. As an alternative, the repository also provides a link to a Docker image.

### Individual BiFC validation in live cells

Without inhibitor: HEK293T were seeded on glass coverslips in 6-well plates (3.5 x 10^5^ cells per well). One day later, a total of 500 ng of plasmid DNA (250 ng of pLIX-VN-ERK and 250 ng of pLIX-CC-ORF) was transfected per well using jetPRIME (ref:114-15, Polyplus Transfection, France) and following manufacturer’s instructions. The cells were simultaneously cotransfected with 100 ng of pLIX-mCherry plasmids to assess transfection efficiency and treated with dox (100 ng/mL final). After 18 h of incubation, the cell-coated coverslip was removed and carefully mounted on a glass slide using a 4% formaldehyde solution for imaging under Zeiss LSM780 confocal microscope (Carl Zeiss, Jena, Germany).

With inhibitor: HEK293T cells were seeded (2.5 x 10^5^ cells) in μDish IBIDI (ref:81156). For U0126 treatment, it was crucial to pre-treat the cells before transfection. One day after plating, cells were pretreated with U0126 (20 µM final) or DMSO (20 µM final) for control. The following day, a total of 600 ng of plasmid DNA (250 ng of pLIX-VN-ERK, 250 ng of pLIX-CC-ORF and 100 ng of pLIX-mCherry) was transfected per well using jetPRIME (ref:114-15, Polyplus Transfection, France) and following manufacturer’s instructions. During transfection, the cells were simultaneously treated with U0126 (30 µM final) or DMSO (30 µM final) and dox (100 ng/mL final). After 24 h of incubation, live cell imaging was performed using a Zeiss LSM780 confocal microscope (Carl Zeiss, Jena, Germany). For EI52, no pre-treatment was required. The same conditions used for U0126 were applied without pre-treating the cells. At the time of transfection, the cells were treated with EI52 (12 µM final) or DMSO (12 µM final) for control.

### splitFAST

HEK293T cells were seeded (2.5 x 10^5^ cells) in μDish IBIDI (ref:81156) coated with poly-L-lysine the day before beginning the experiment. Once the cells reached approximately 80% confluence, they were transfected with splitFAST2 constructs (1 µg of pLIX-NFAST-ERK1 and 1 µg of pLIX-CFAST-ORF) using jetPRIME (Ref 114-15, Polyplus Transfection, France) according to the manufacturer’s recommendations. As a control, cells were simultaneously transfected with BiFC constructs (200 ng of pLIX-VN-ERK1 and 200 ng of pLIX-CC-ORF) from the same partner pair. To study the dissociation of the splitFAST2 system, cells were treated with dox (500 ng/mL final) at the time of transfection. 12 h after, the medium was replaced with complete medium without phenol red to remove dox and phenol red (to avoid fluorescence interference with the HBR signal) and the cells were subsequently treated with EI52 (12 µM final) or U0126 (10 µM final) or DMSO (12 µM or 10 µM final) for control. About 6h after treatment, cells were imaged live under Zeiss LSM780 confocal microscope (Carl Zeiss, Jena, Germany). The optimal HBR fluorogen concentration required to visualize the association or dissociation of the splitFAST2 system was determined for each interaction tested. A 100 µL solution containing the appropriate concentration of fluorogen was added live just before doing confocal acquisitions.

### Clustering interactome and functional enrichment analysis

Functional networks of ERK interactome were generated with STRING software (https://string-db.org/), based on 0.15 interaction score of experimental evidence, database and pathway co-occurrence. Visualization of networks was built with Cytoscape free software conserving the default parameters and clustered according to their biological functional enrichment (https://cytoscape.org/). The GO-Term annotations and over-represented GO-Term related to biological process analysis were performed with Metascape (https://metascape.org/gp/index.html#/main/step1).

## Acknowledgements

We thank Lucas Pons for his help in doing BiFC and splitFAST2 experiments, and Christelle Forcet for her careful reading of the manuscript. This project was supported by funding from Ecole Normale Supérieure de Lyon (ENSL), La Ligue Nationale Contre le Cancer (OPE-2020-0024), Projet Fondation ARC (PJA 20191209567), Pack Ambition Région Rhône-Alpes-Auvergne 2020 and Institut National du Cancer PLBIO (TR-2020-119).

## Supplementary Table legends

**Supplementary Table 1**

List of integrated CC-ORFs in the BiFC-ERKeome cell line

**Supplementary Table 2**

Log2FC enrichment and p-values for each CC-ORF with VN-ERK1 when compared to VN and calculated by considering the three biological replicates.

**Supplementary Table 3**

List of the 20 selected interactions with their corresponding Log2FC enrichment and adjusted p-values.

**Supplementary Table 4**

Raw values of Log2FC enrichment and p-values for each CC-ORF with VN-ERK1 when tested in the presence of U0126 and calculated by considering the three biological replicates. The 24 U0126-sensitive interactions that met the selection criteria are highlighted. Positive ERK1 interactions found in the absence of drug are indicated with an asterisk.

**Supplementary Table 5**

Raw values of Log2FC enrichment and p-values for each CC-ORF with VN-ERK1 when tested in the presence of EI52 and calculated by considering the three biological replicates. The 25 EI52-sensitive interactions that met the selection criteria are highlighted. Positive ERK1 interactions found in the absence of drug are indicated with an asterisk.

**Supplementary Table 6**

Raw data from the three replicates of the BiFC-ERKeome screen with VN-ERK1 or VN, in wild type condition or in the presence of U0126 or EI52.

---

**Supplementary Figure 1.**
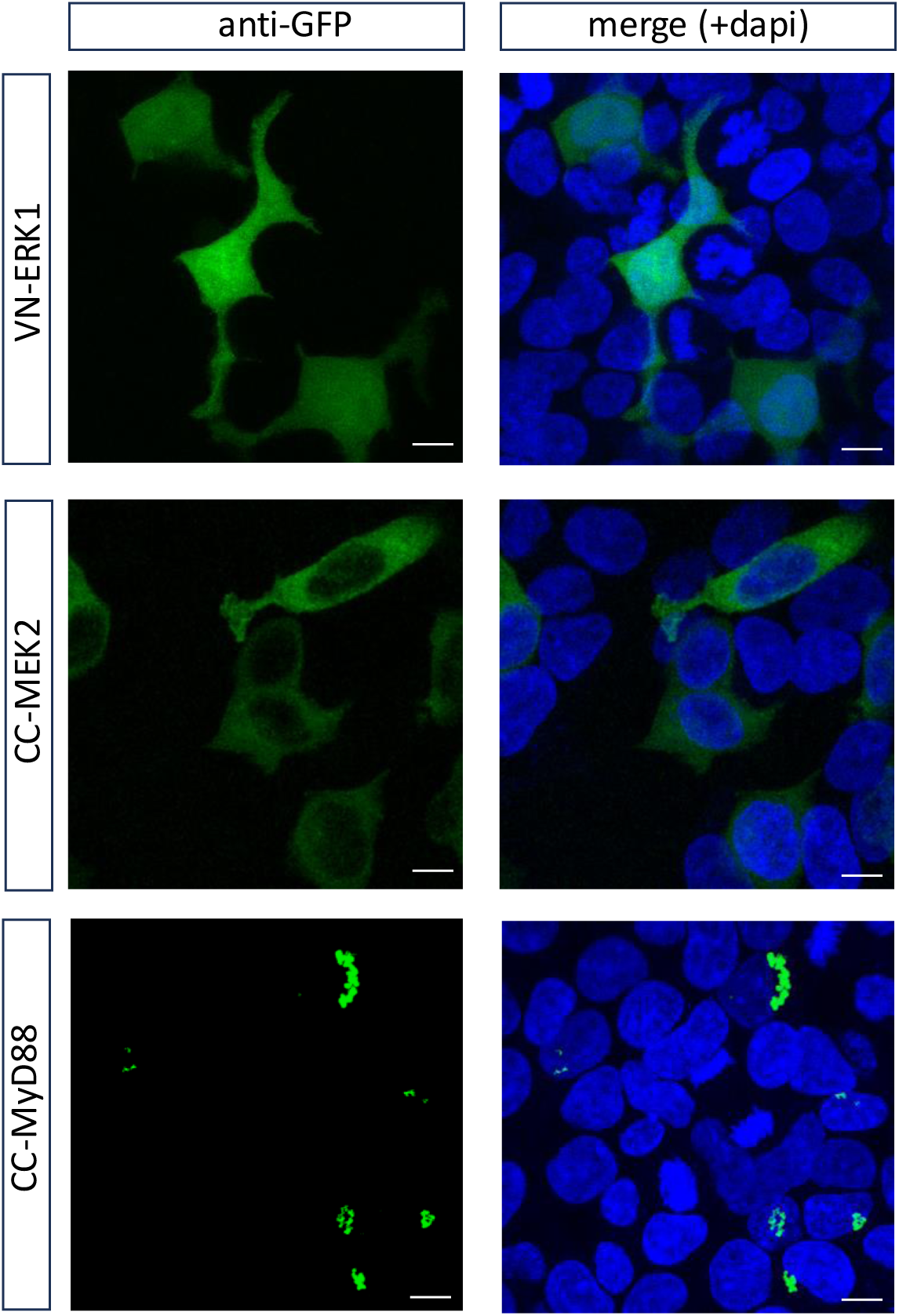
Fluorescent immunostaining of the VN-ERK1, CC-MEK2 and CC-MyD88 constructs transfected in HEK293T cells and revealed with an anti-GFP (green) recognizing the VN and CC fragment, as indicated. Dapi (blue) stains for nuclei. Scale bar=10 µm.

**Supplementary Figure 2.**
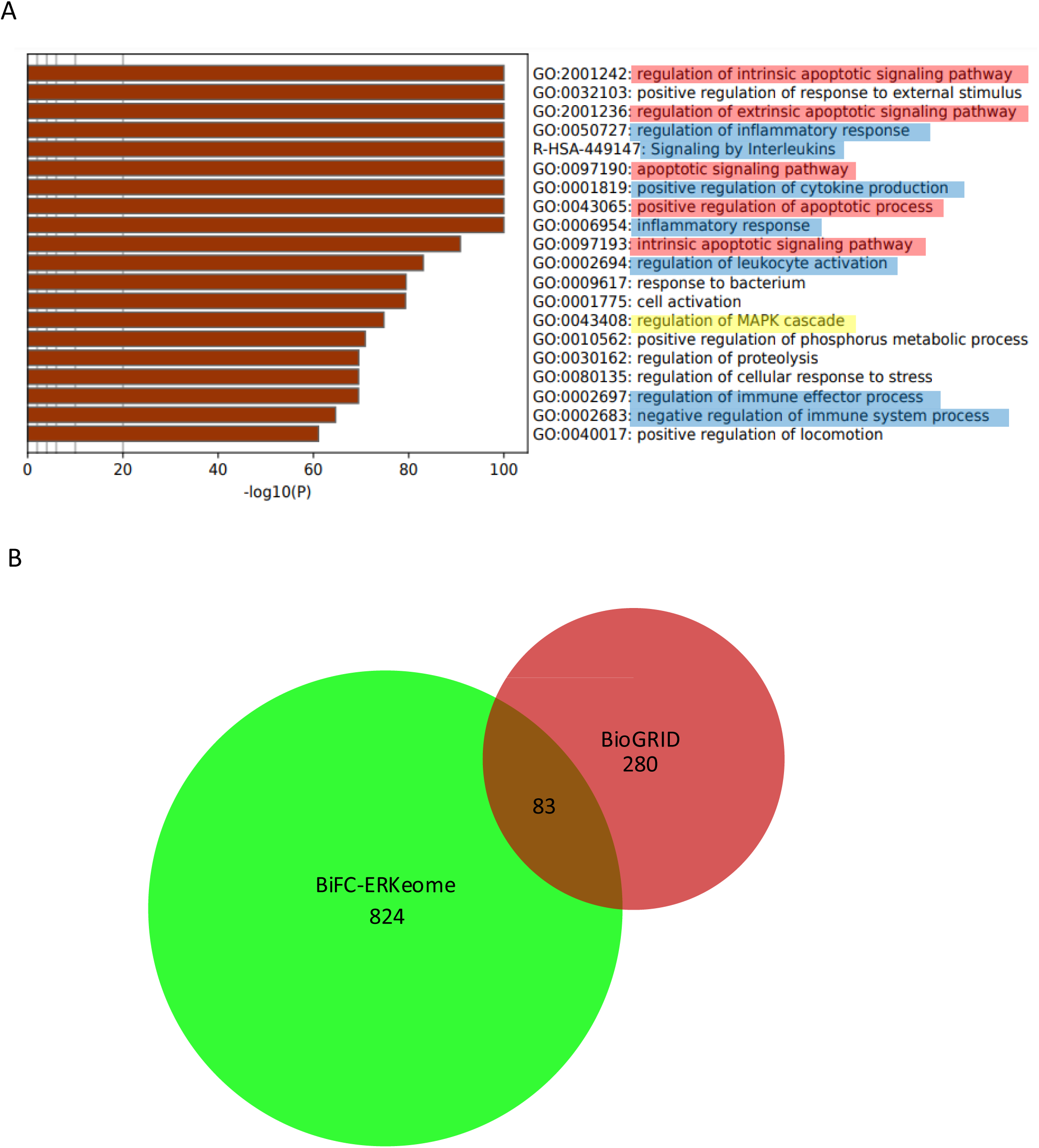
**A.** GO term enrichment of the integrated BiFC-ERKeome in HEK293T cells. Color code indicates functionally related GO terms (apoptosis in red, MAPK pathway in yellow, immunity in blue). **B.** Venn diagram showing the overlap between the integrated BiFC-ERKeome and the know ERK1 interactome (as provided in the BioGrid database).

**Supplementary Figure 3.**
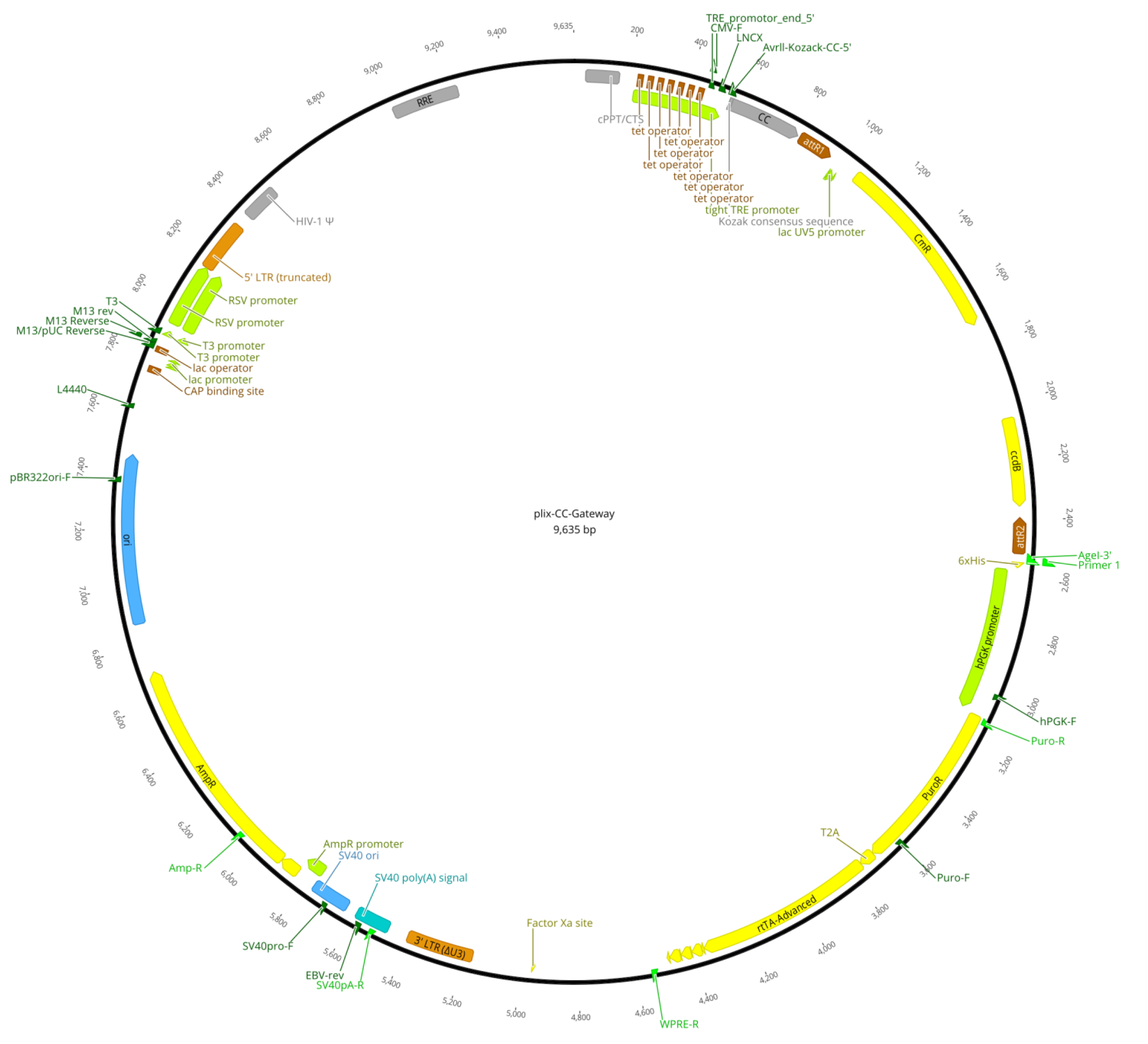
Map of the pLix-CC-GW vector used to clone each candidate ORF in fusion with the C-terminal fragment of Cerulean (CC). GW: Gateway.

**Supplementary Figure 4.**
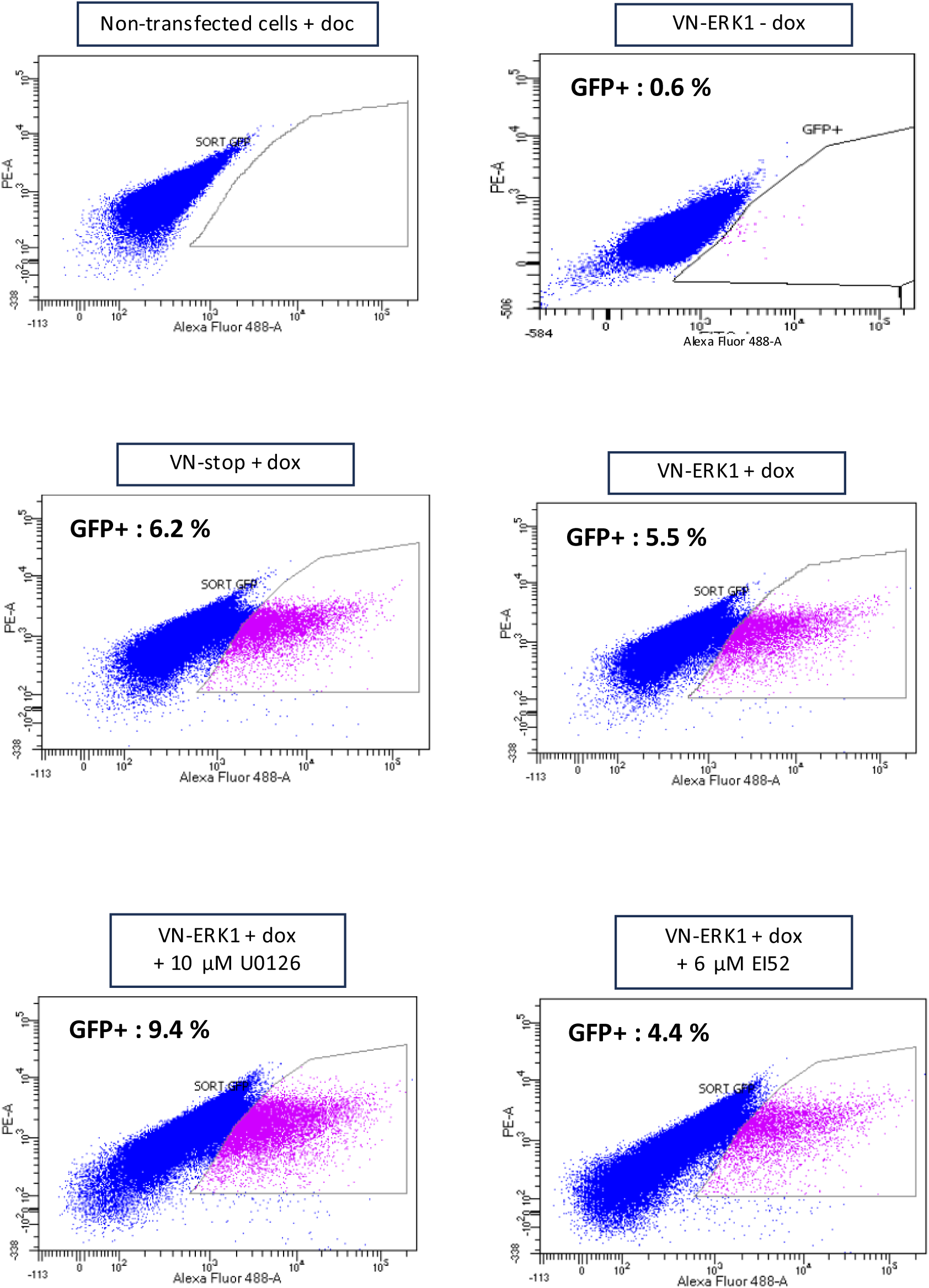
Graphs showing the proportion of fluorescent cells when analyzed by the cell sorter in each condition, as indicated. The gate-filter used to specifically sort the fluorescent cells is indicated (black line).

**Supplementary Figure 5.**
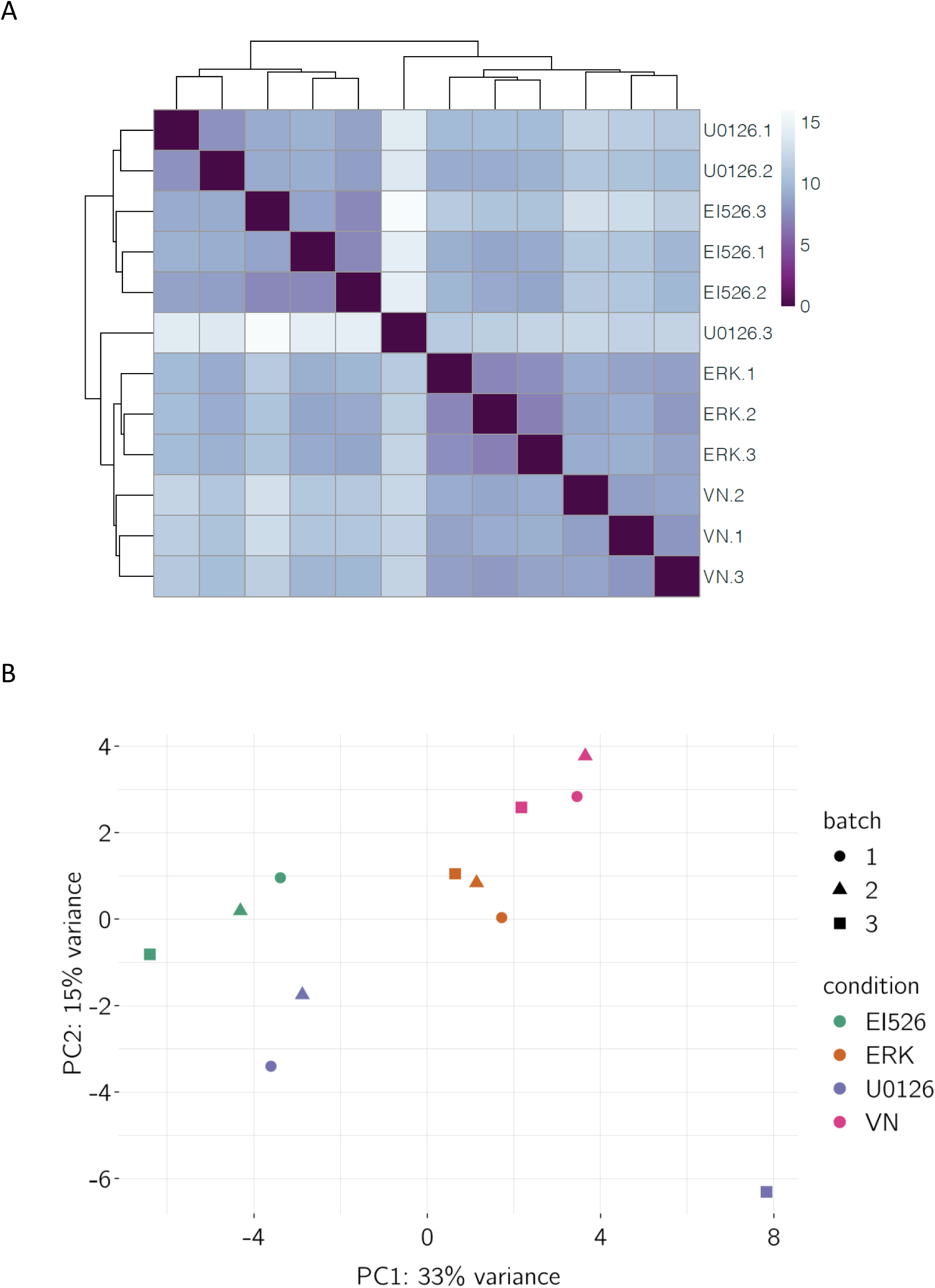
Clustering of the different conditions. **A.** Hierarchical clustering of the different conditions and for the three biological replicates, as indicated. **B.** PCA of the different conditions (indicated by different colors) and for the three biological replicates (batch 1, 2 or 3).

**Supplementary Figure 6.**
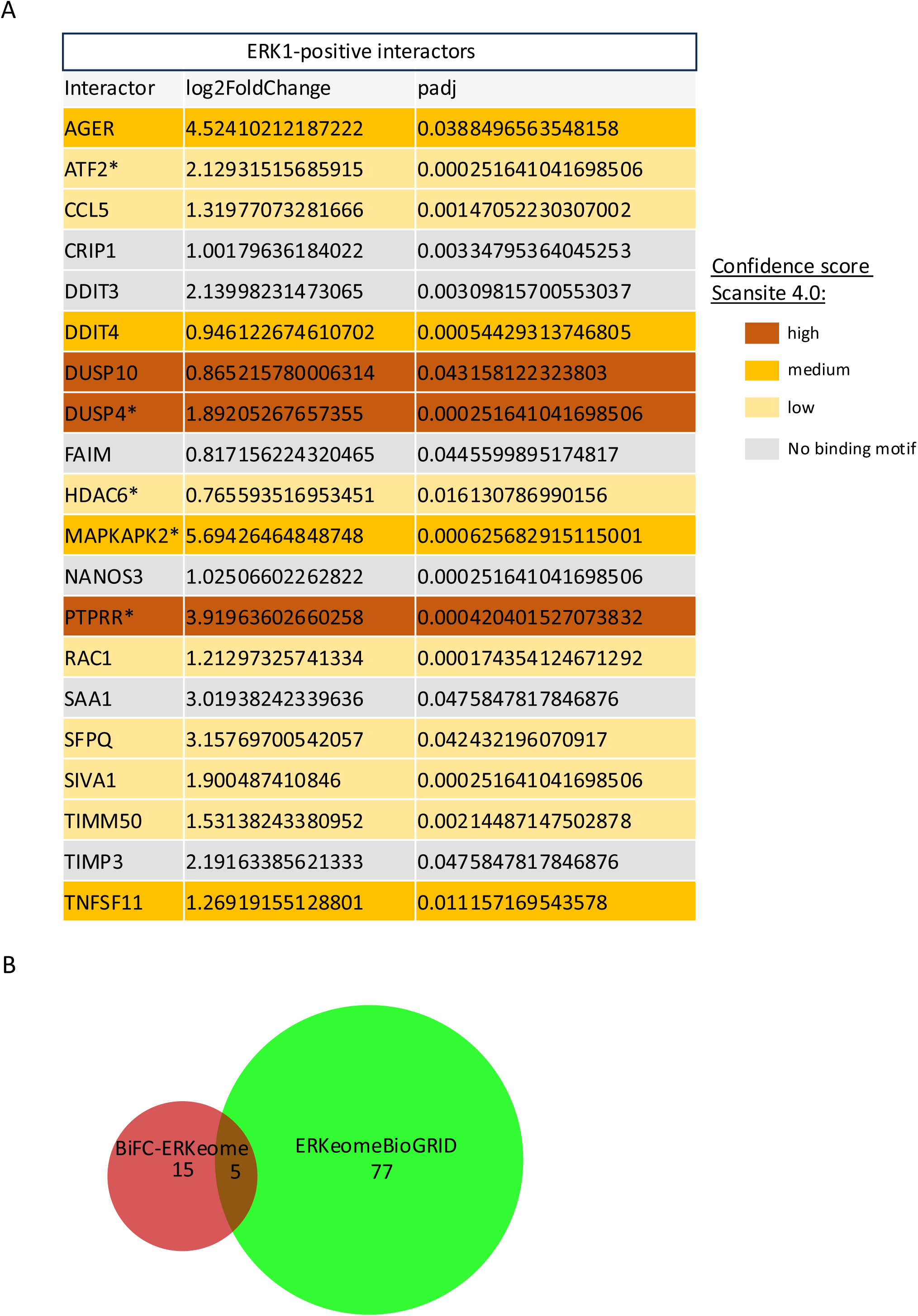
**A.** List of the 20 selected ERK1 interactors after applying the log2FC and adjusted p-values filtering. Each interactor is color-filled depending on its confidence score for containing ERK D-binding domain, as indicated. Interactors with an asterisk are known ERK1 cofactors. **B.** Venn diagram showing the overlap between BiFC-positive interactions and the known cofactors of ERK1 in the BiFC-ERKeome (as provided in the BioGrid database).

**Supplementary Video 1.**
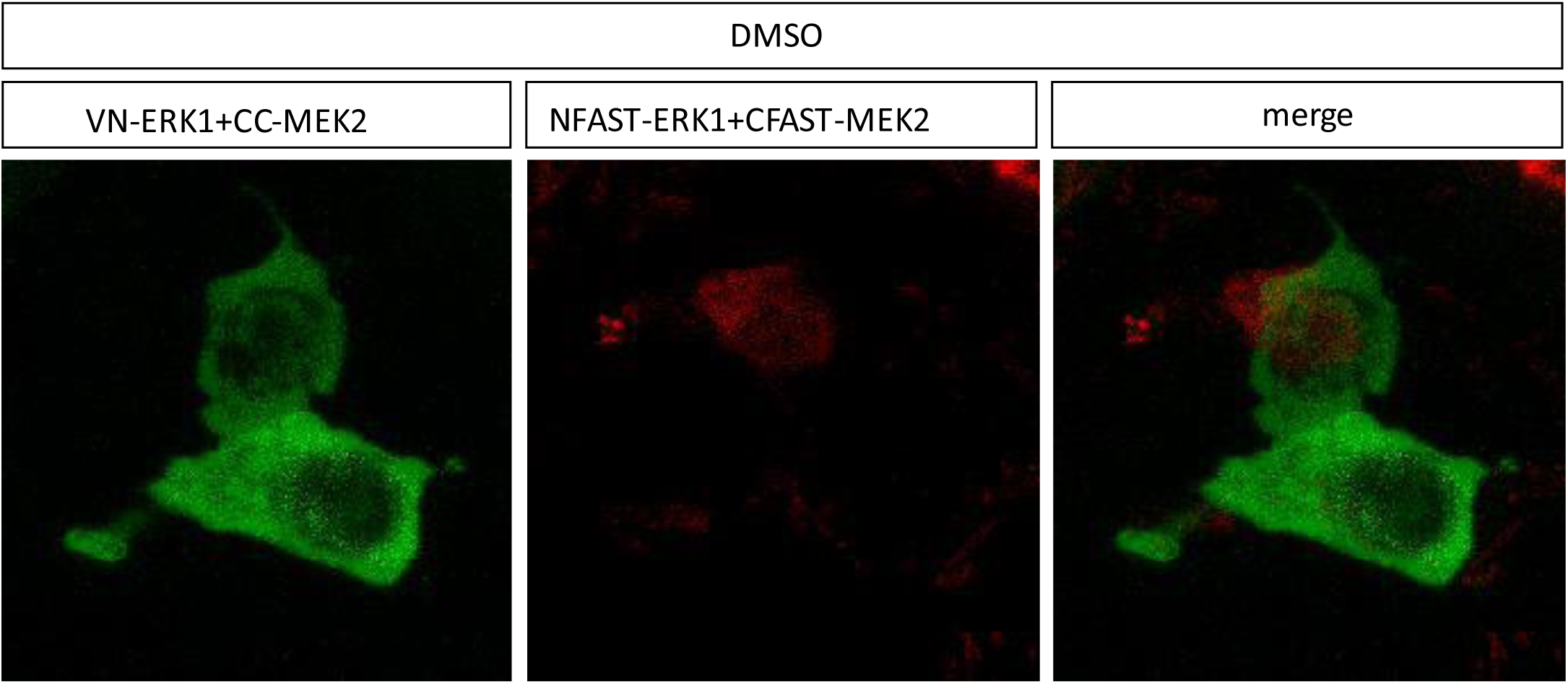
BiFC of ERK1/MEK2 interaction in the presence of DMSO from 0 to 10 minutes after the addition of the HBR ligand. DMSO that was added 12h post-transfection and cells were cultured for additional 6h in the presence of DMSO before live imaging.

**Supplementary Video 2.**
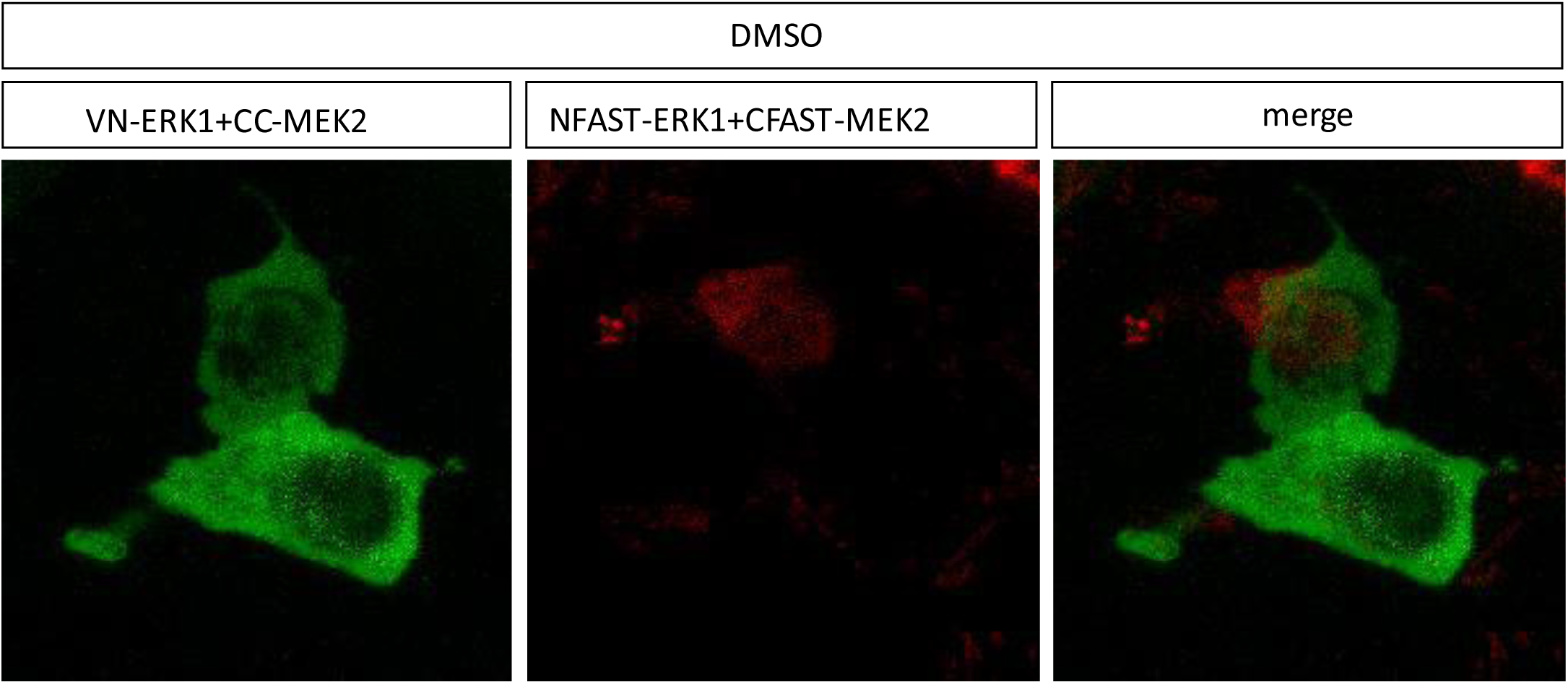
splitFAST2 of ERK1/MEK2 interaction in the presence of DMSO from 0 to 10 minutes after the addition of the HBR ligand. DMSO that was added 12h post-transfection and cells were cultured for additional 6h in the presence of DMSO before live imaging.

**Supplementary Video 3.**
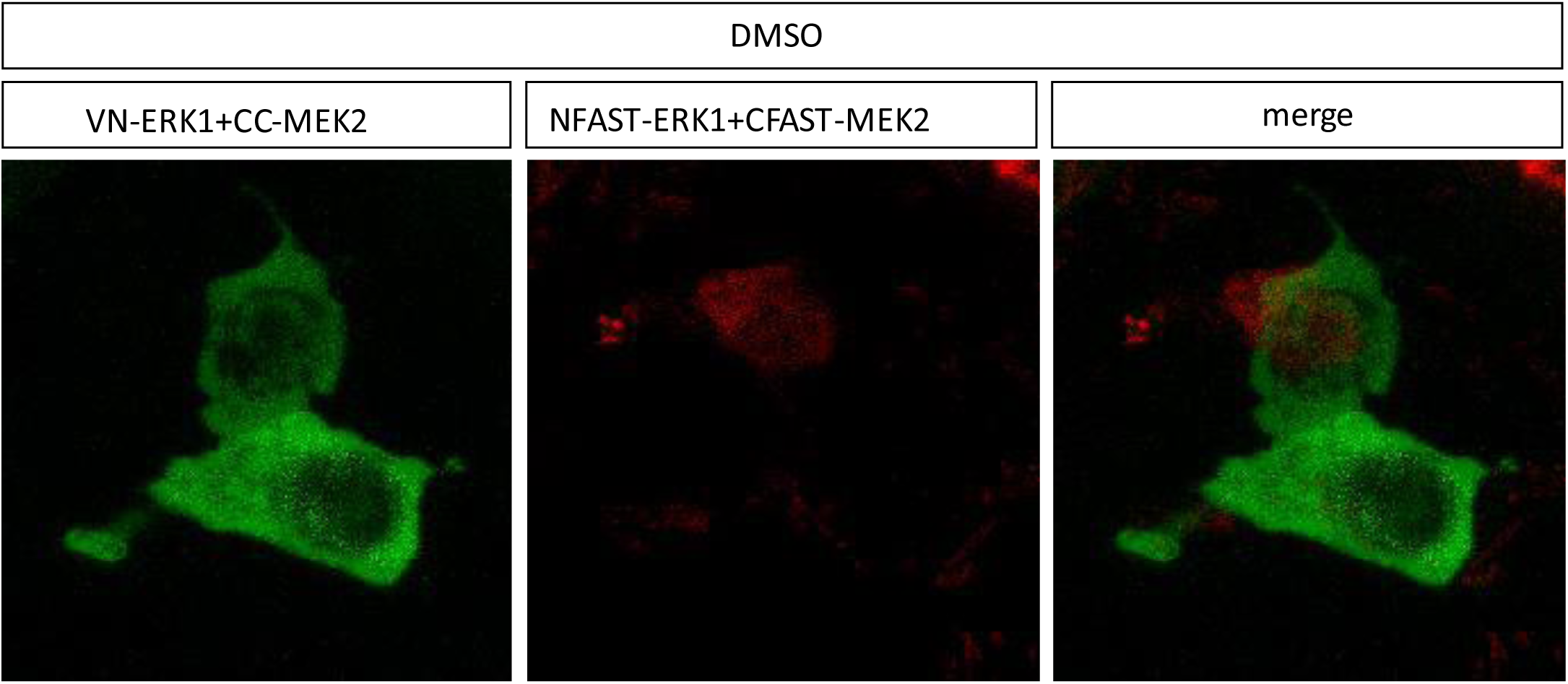
BiFC and splitFAST2 of ERK1/MEK2 interaction in the presence of DMSO from 0 to 10 minutes after the addition of the HBR ligand. DMSO that was added 12h post-transfection and cells were cultured for additional 6h in the presence of DMSO before live imaging.

**Supplementary Video 4.**
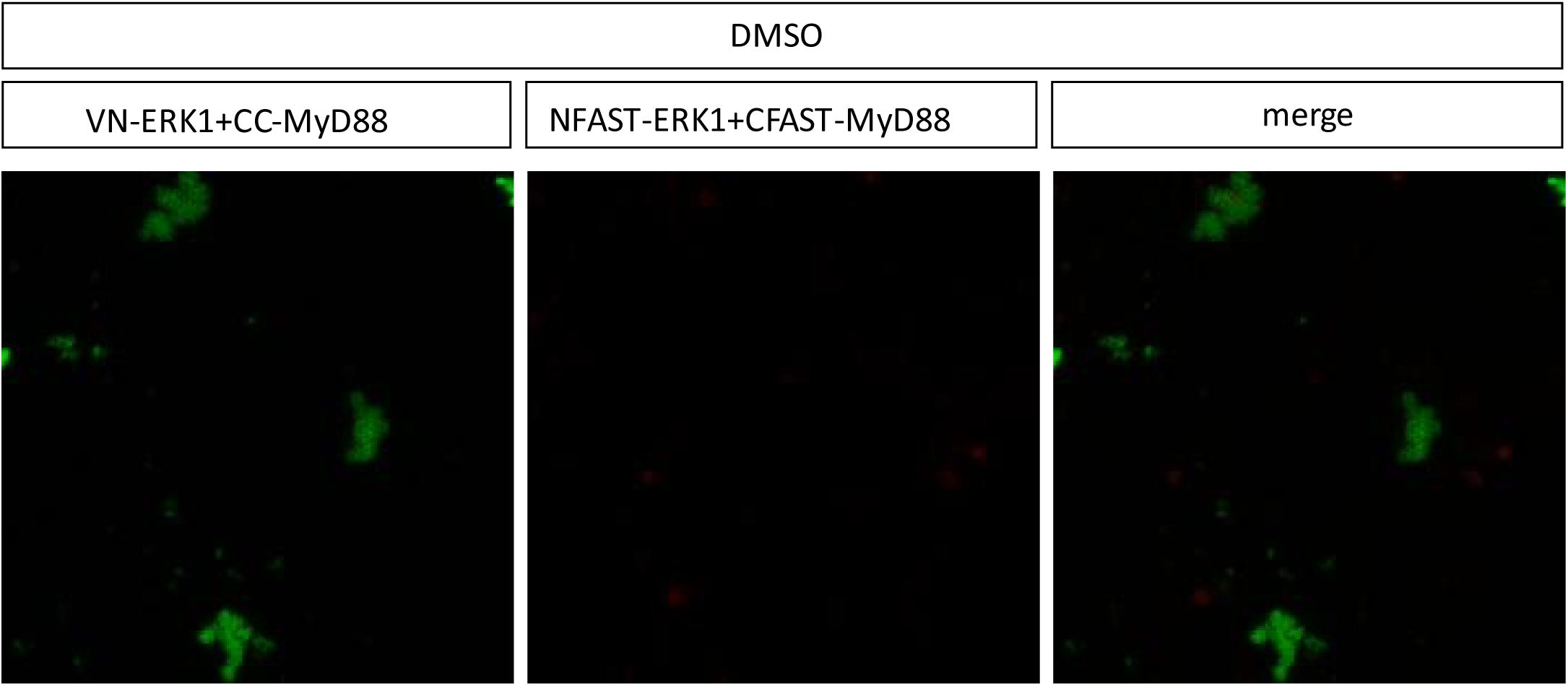
BiFC of ERK1/MyD88 interaction in the presence of DMSO from 0 to 10 minutes after the addition of the HBR ligand. DMSO that was added 12h post-transfection and cells were cultured for additional 6h in the presence of DMSO before live imaging.

**Supplementary Video 5.**
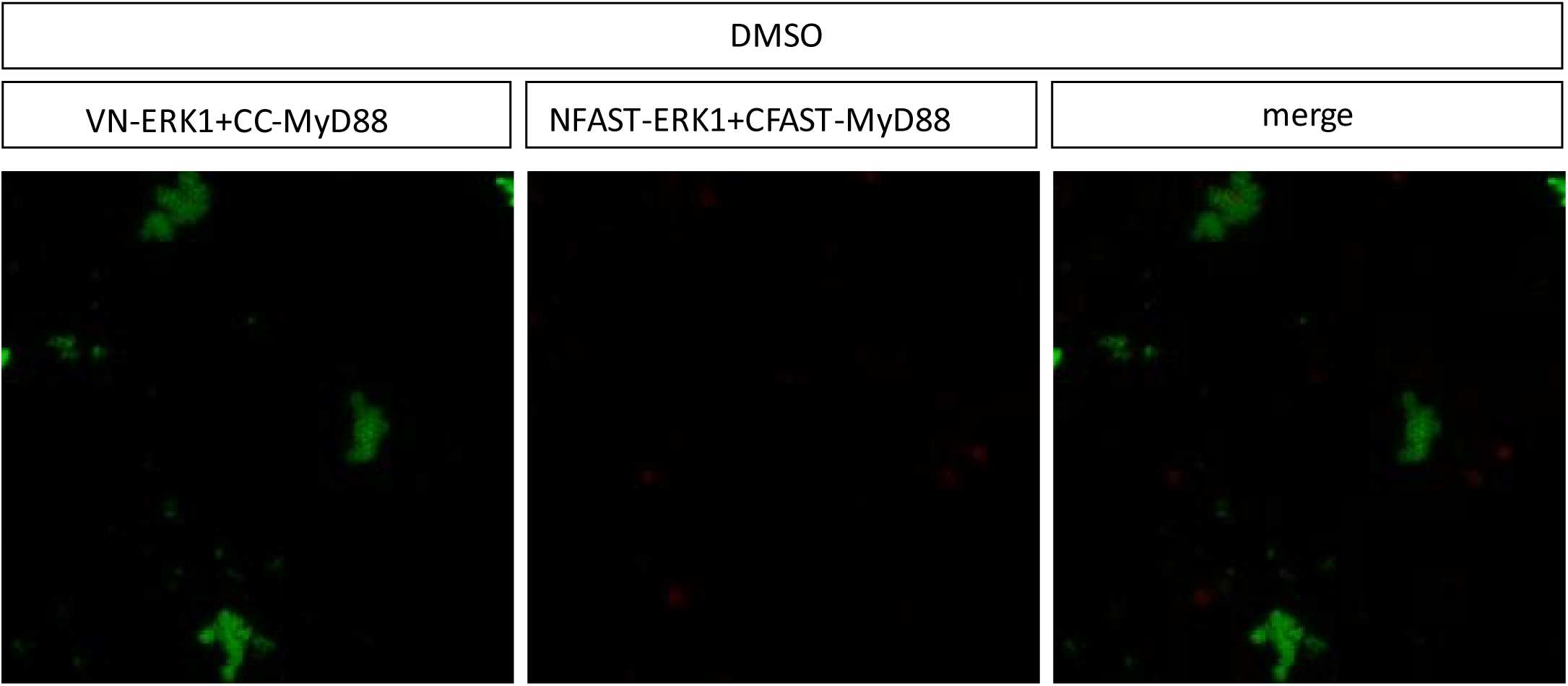
splitFAST2 of ERK1/MyD88 interaction in the presence of DMSO from 0 to 10 minutes after the addition of the HBR ligand. DMSO that was added 12h post-transfection and cells were cultured for additional 6h in the presence of DMSO before live imaging.

**Supplementary Video 6.**
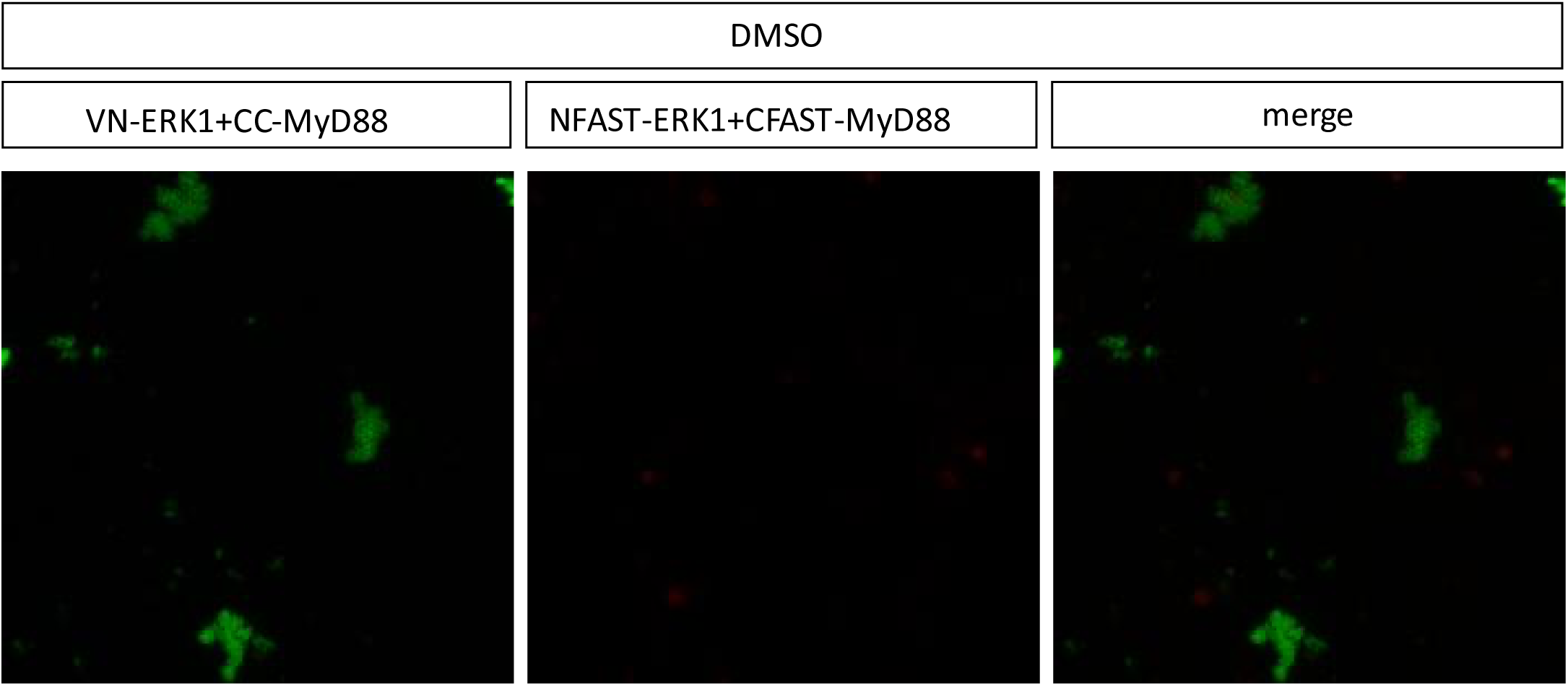
BiFC and splitFAST2 of ERK1/MyD88 interaction in the presence of DMSO from 0 to 10 minutes after the addition of the HBR ligand. DMSO that was added 12h post-transfection and cells were cultured for additional 6h in the presence of DMSO before live imaging.

**Supplementary Video 7.**
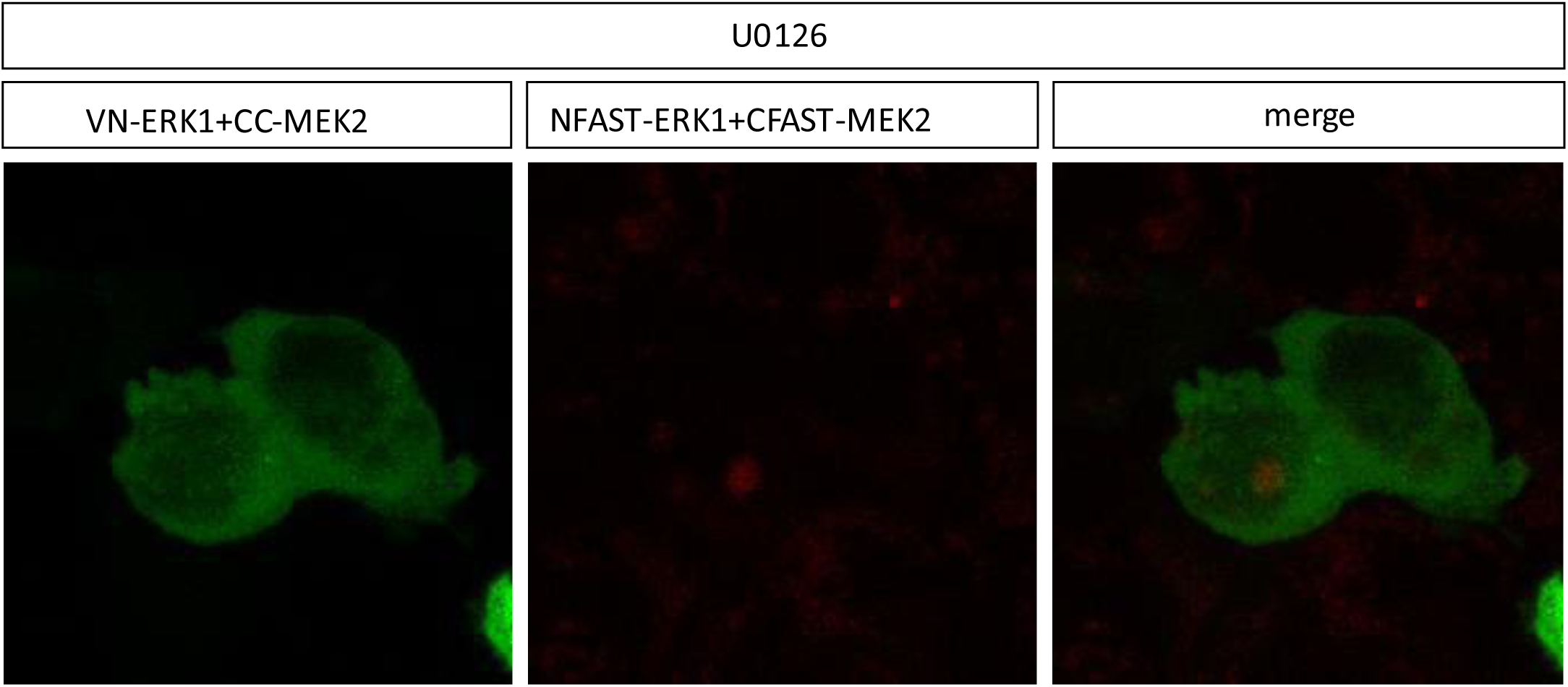
BiFC of ERK1/MEK2 interaction in the presence of DMSO+U0126 from 0 to 10 minutes after the addition of the HBR ligand. DMSO+U0126 that was added 12h post-transfection and cells were cultured for additional 6h in the presence of DMSO+U0126 before live imaging.

**Supplementary Video 8.**
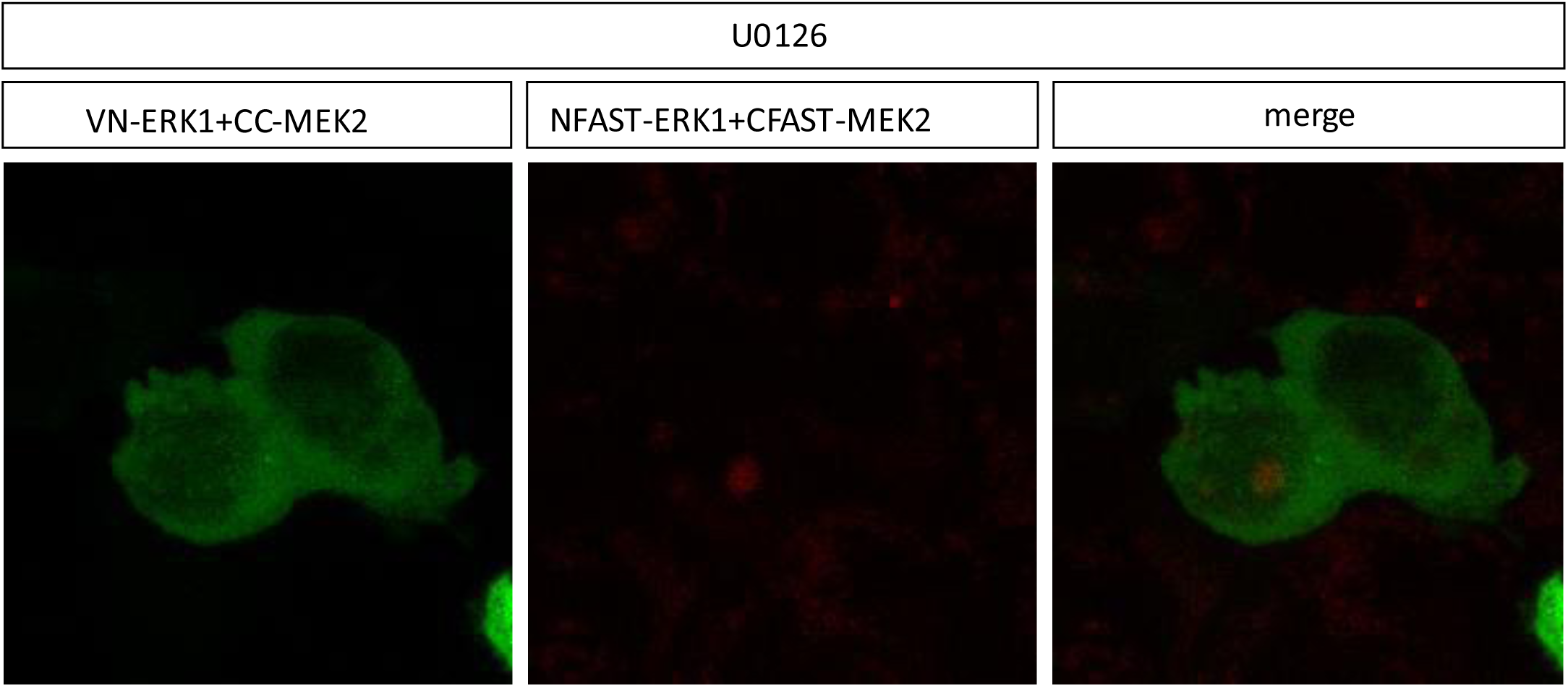
splitFAST2 of ERK1/MEK2 interaction in the presence of DMSO+U0126 from 0 to 10 minutes after the addition of the HBR ligand. DMSO+U0126 that was added 12h post-transfection and cells were cultured for additional 6h in the presence of DMSO+U0126 before live imaging.

**Supplementary Video 9.**
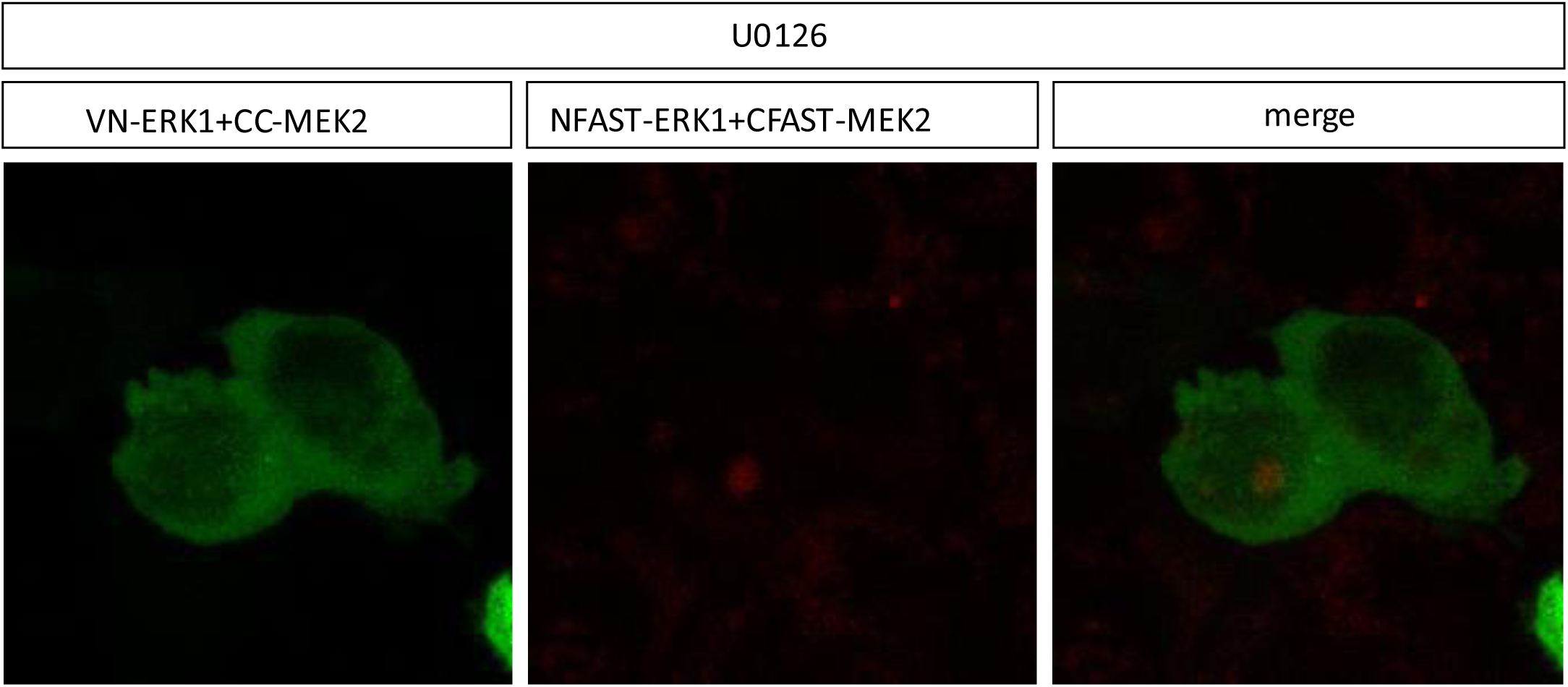
BiFC and splitFAST2 of ERK1/MEK2 interaction in the presence of DMSO+U0126 from 0 to 10 minutes after the addition of the HBR ligand. DMSO+U0126 that was added 12h post-transfection and cells were cultured for additional 6h in the presence of DMSO+U0126 before live imaging.

**Supplementary Video 10.**
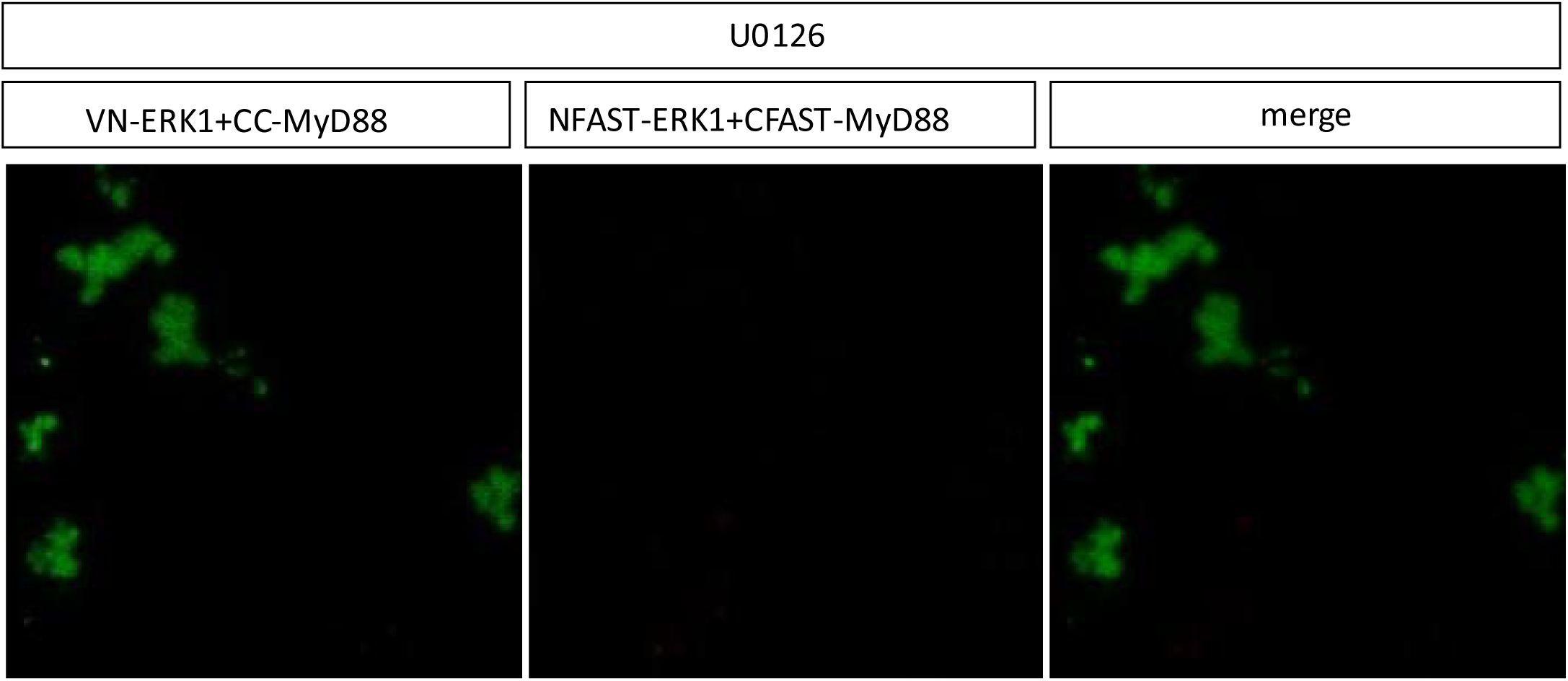
BiFC of ERK1/MyD88 interaction in the presence of DMSO+U0126 from 0 to 10 minutes after the addition of the HBR ligand. DMSO+U0126 that was added 12h post-transfection and cells were cultured for additional 6h in the presence of DMSO+U0126 before live imaging.

**Supplementary Video 11.**
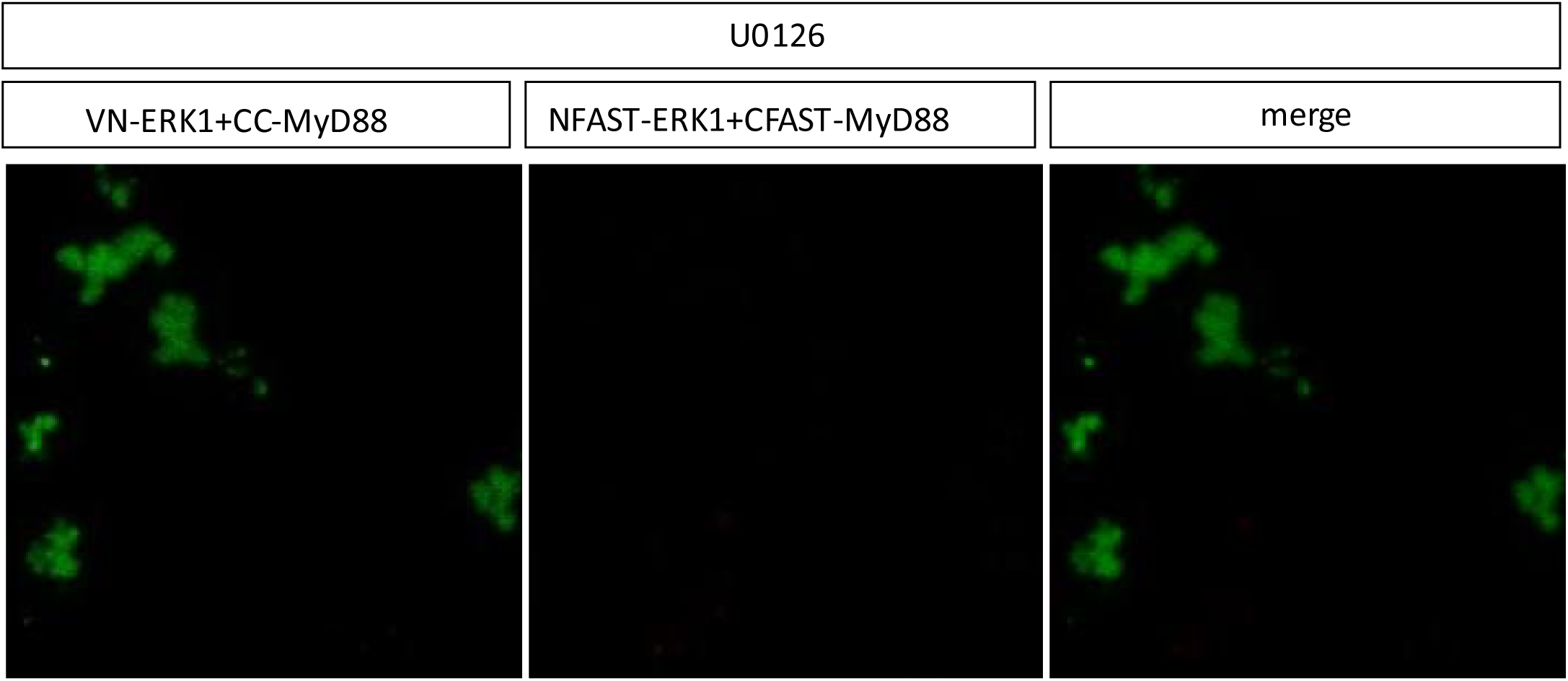
splitFAST2 of ERK1/MyD88 interaction in the presence of DMSO+U0126 from 0 to 10 minutes after the addition of the HBR ligand. DMSO+U0126 that was added 12h post-transfection and cells were cultured for additional 6h in the presence of DMSO+U0126 before live imaging.

**Supplementary Video 12.**
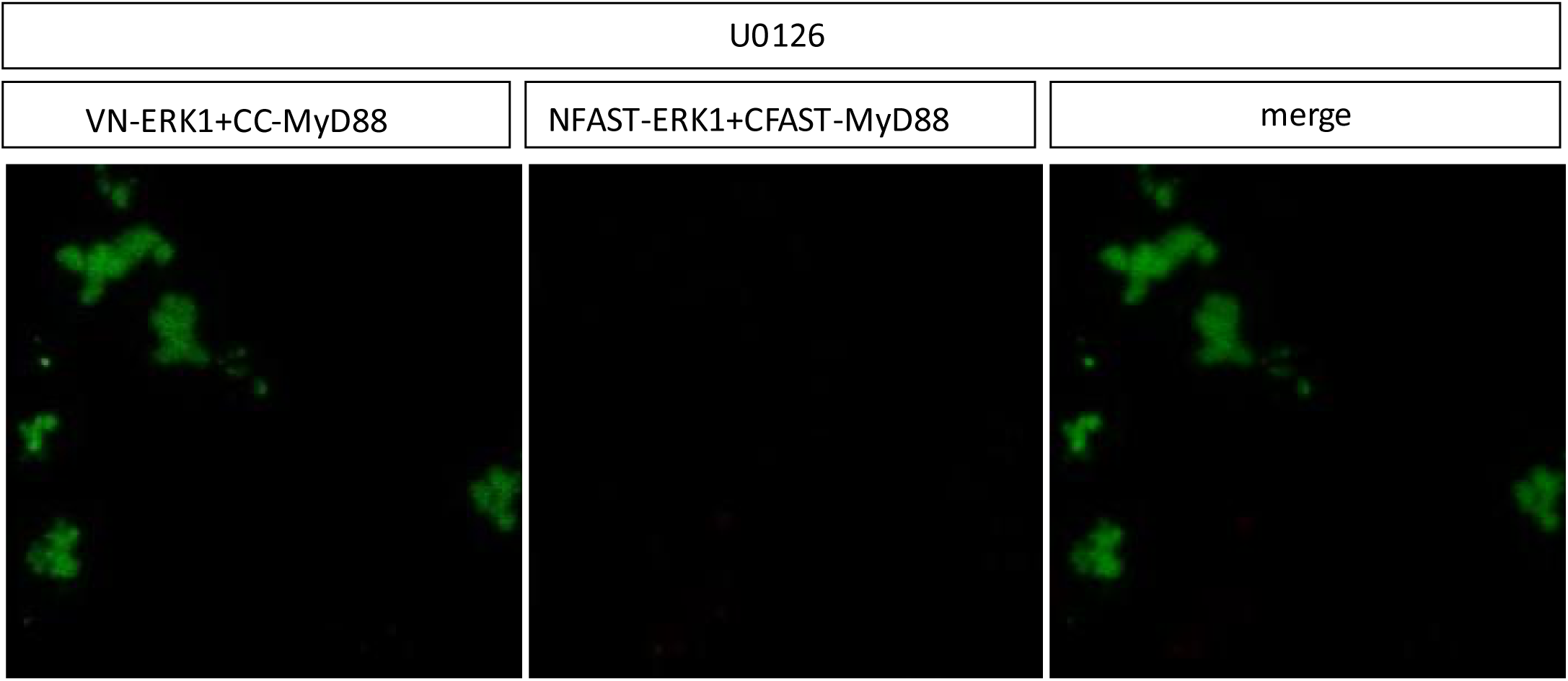
BiFC and splitFAST2 of ERK1/MyD88 interaction in the presence of DMSO+U0126 from 0 to 10 minutes after the addition of the HBR ligand. DMSO+U0126 that was added 12h post-transfection and cells were cultured for additional 6h in the presence of DMSO+U0126 before live imaging.

**Supplementary Video 13.**
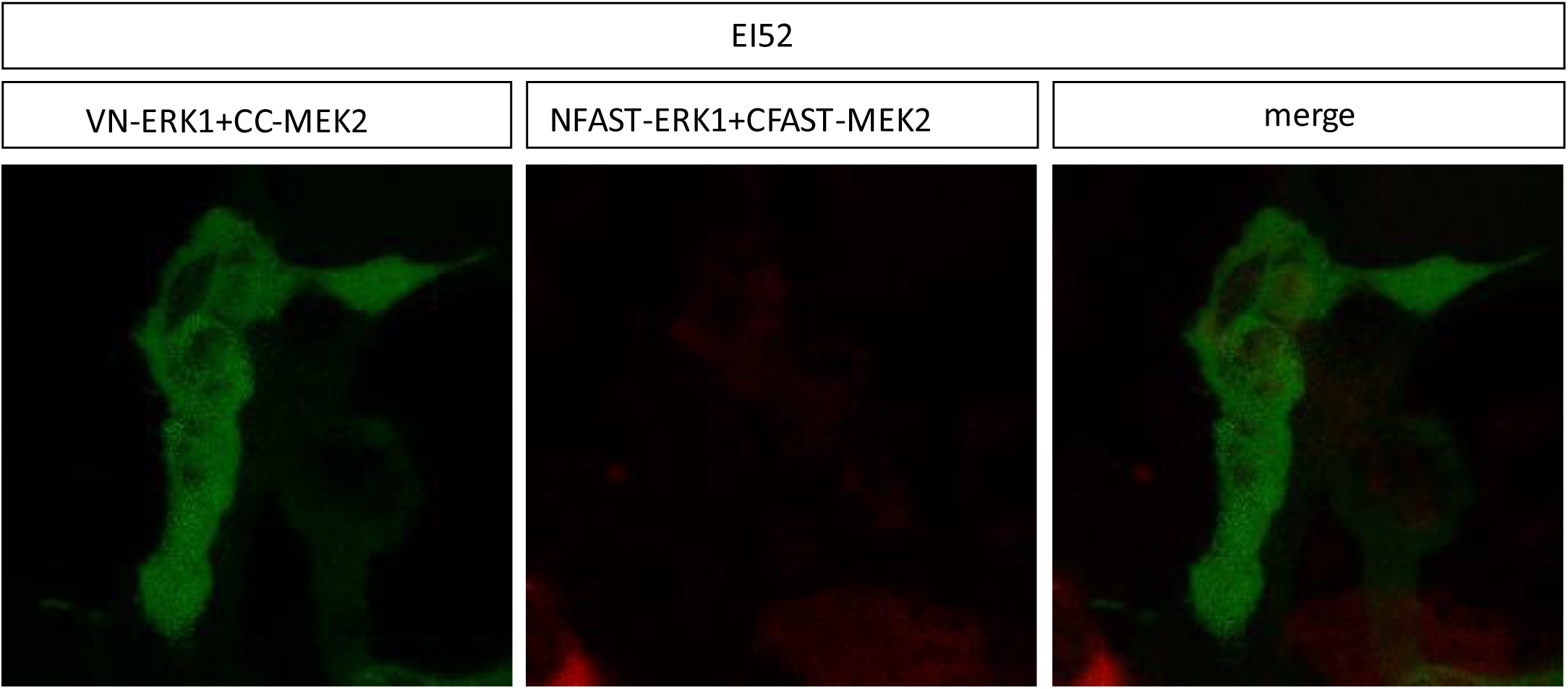
BiFC of ERK1/MEK2 interaction in the presence of DMSO+EI52 from 0 to 10 minutes after the addition of the HBR ligand. DMSO+EI52 that was added 12h post-transfection and cells were cultured for additional 6h in the presence of DMSO+EI52 before live imaging.

**Supplementary Video 14.**
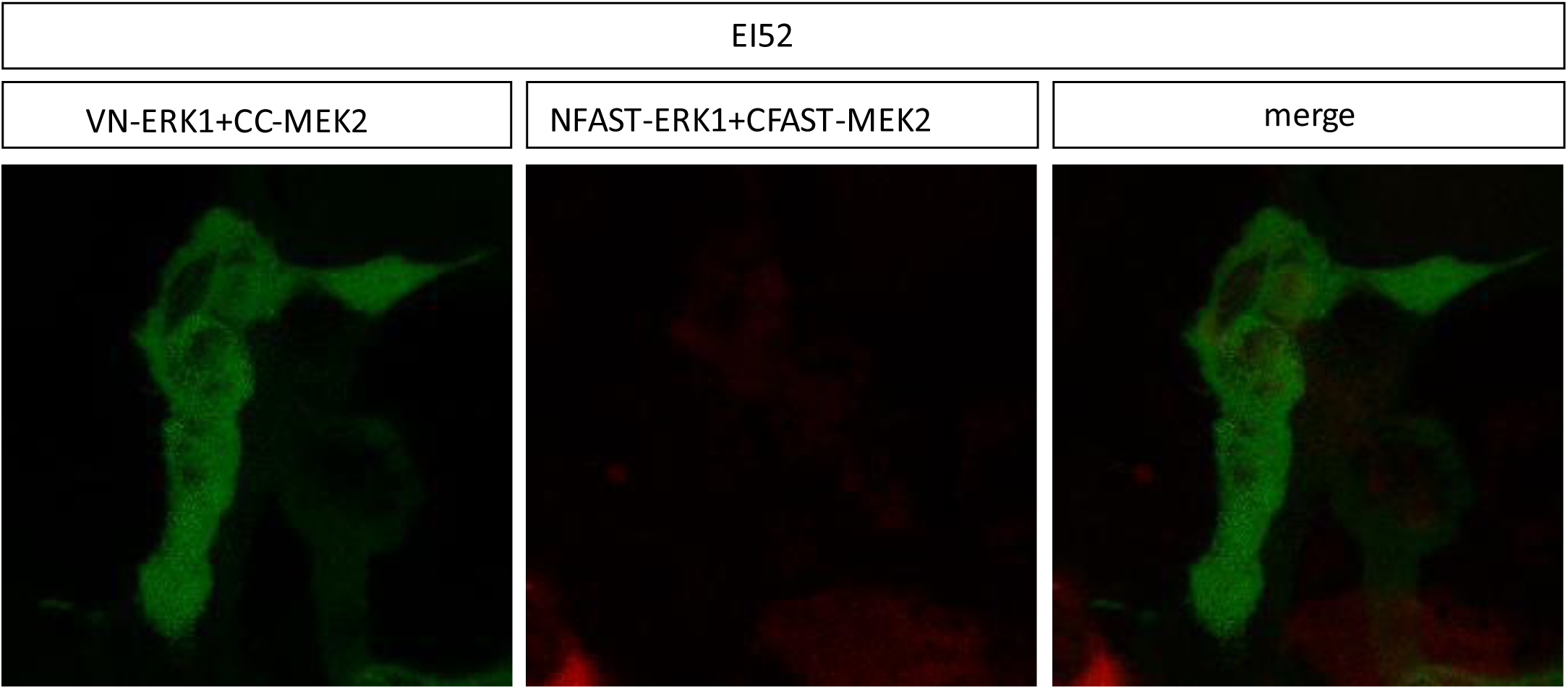
splitFAST2 of ERK1/MEK2 interaction in the presence of DMSO+EI52 from 0 to 10 minutes after the addition of the HBR ligand. DMSO+EI52 that was added 12h post-transfection and cells were cultured for additional 6h in the presence of DMSO+EI52 before live imaging.

**Supplementary Video 15.**
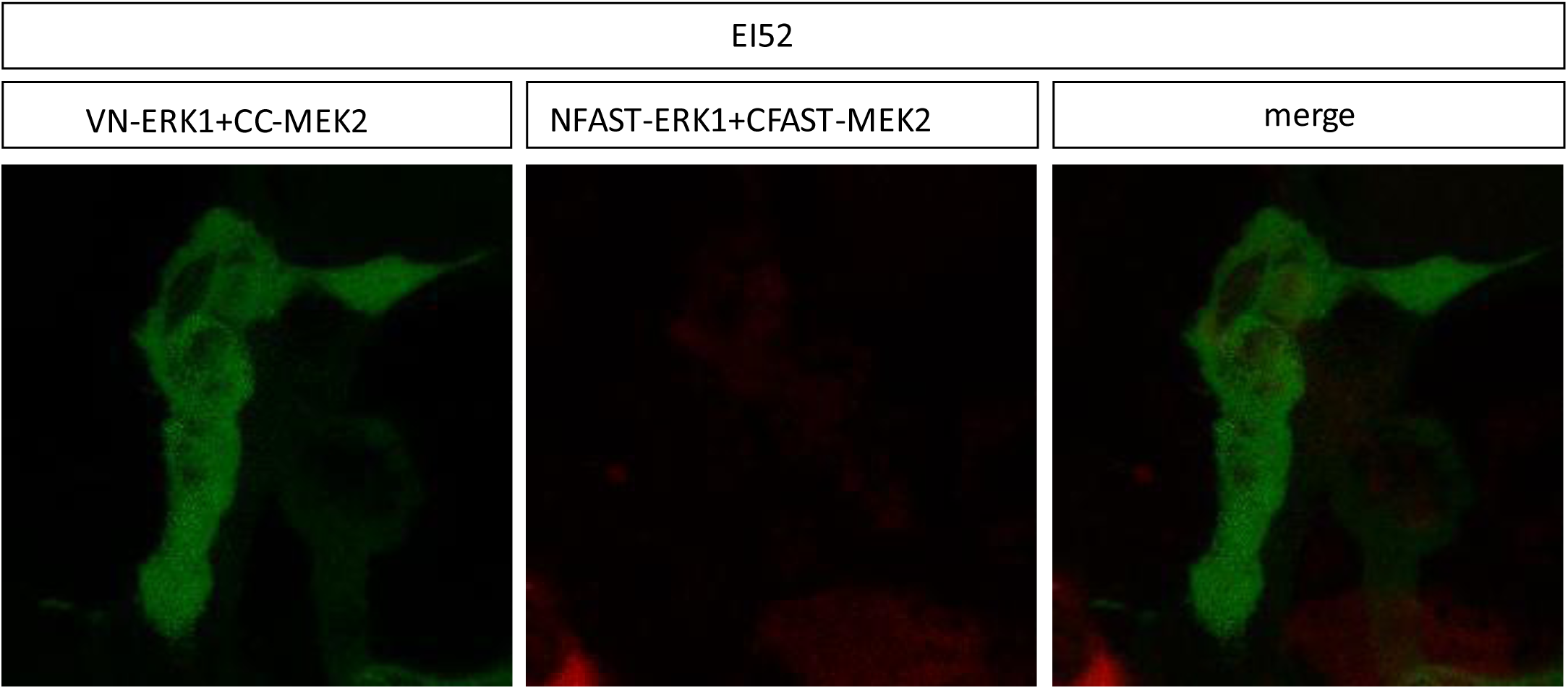
BiFC and splitFAST2 of ERK1/MEK2 interaction in the presence of DMSO+EI52 from 0 to 10 minutes after the addition of the HBR ligand. DMSO+EI52 that was added 12h post-transfection and cells were cultured for additional 6h in the presence of DMSO+EI52 before live imaging.

**Supplementary Video 16.**
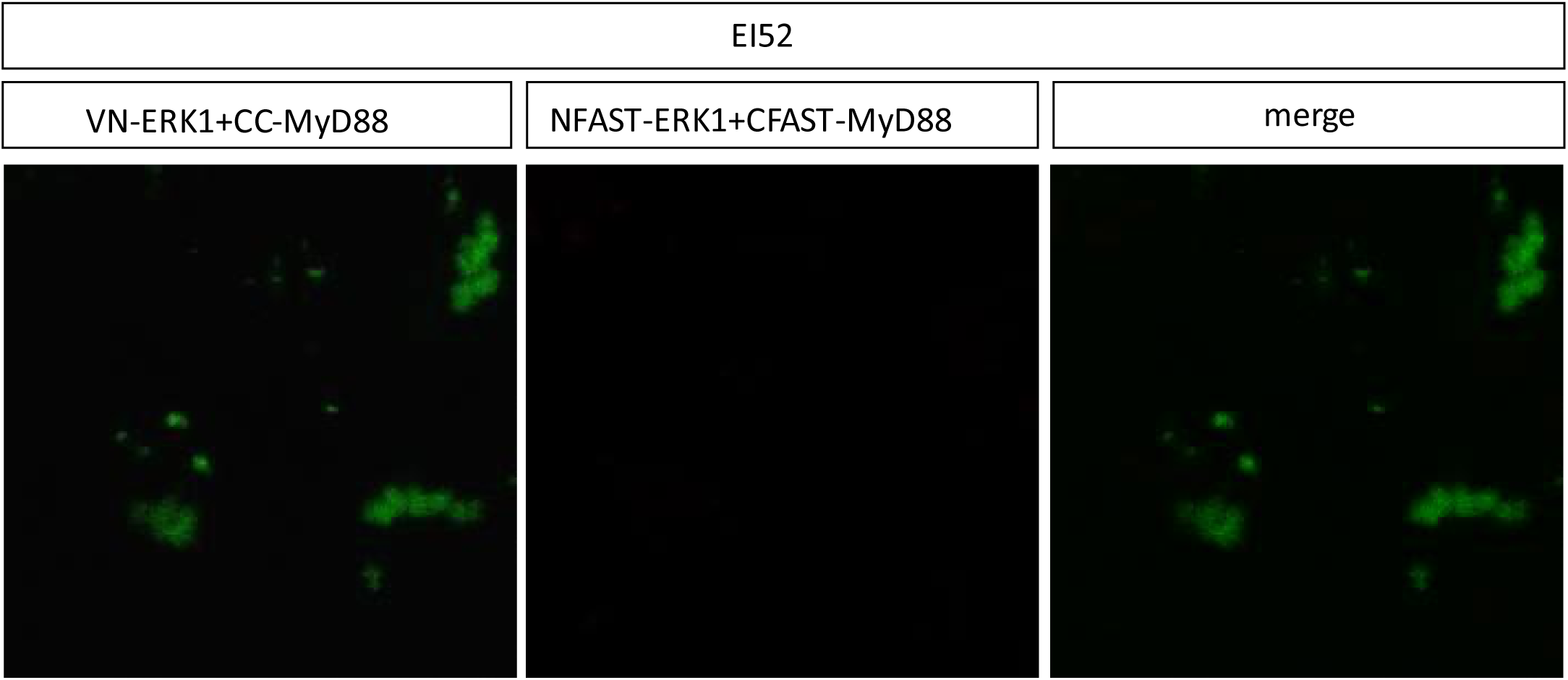
BiFC of ERK1/MyD88 interaction in the presence of DMSO+EI52 from 0 to 10 minutes after the addition of the HBR ligand. DMSO+EI52 that was added 12h post-transfection and cells were cultured for additional 6h in the presence of DMSO+EI52 before live imaging.

**Supplementary Video 17.**
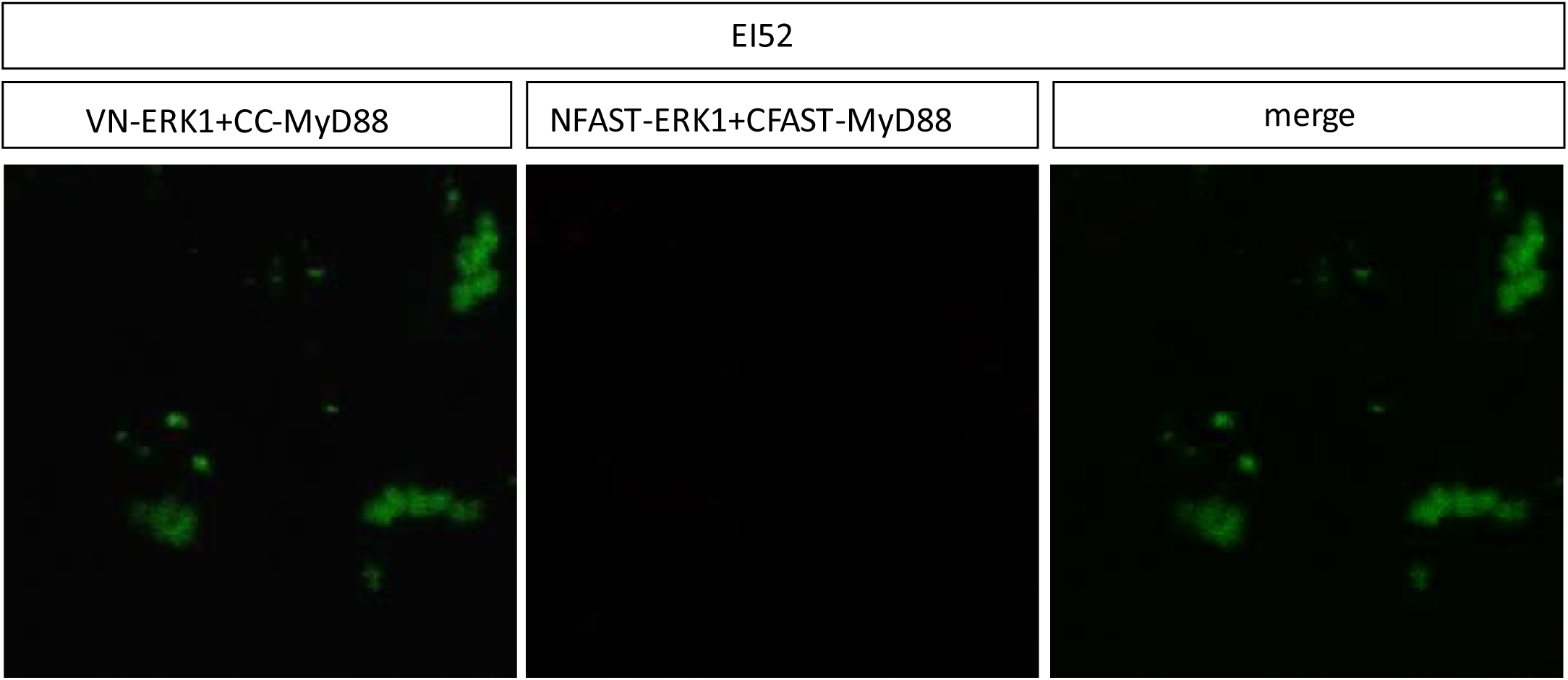
splitFAST2 of ERK1/MyD88 interaction in the presence of DMSO+EI52 from 0 to 10 minutes after the addition of the HBR ligand. DMSO+EI52 that was added 12h post-transfection and cells were cultured for additional 6h in the presence of DMSO+EI52 before live imaging.

**Supplementary Video 18.**
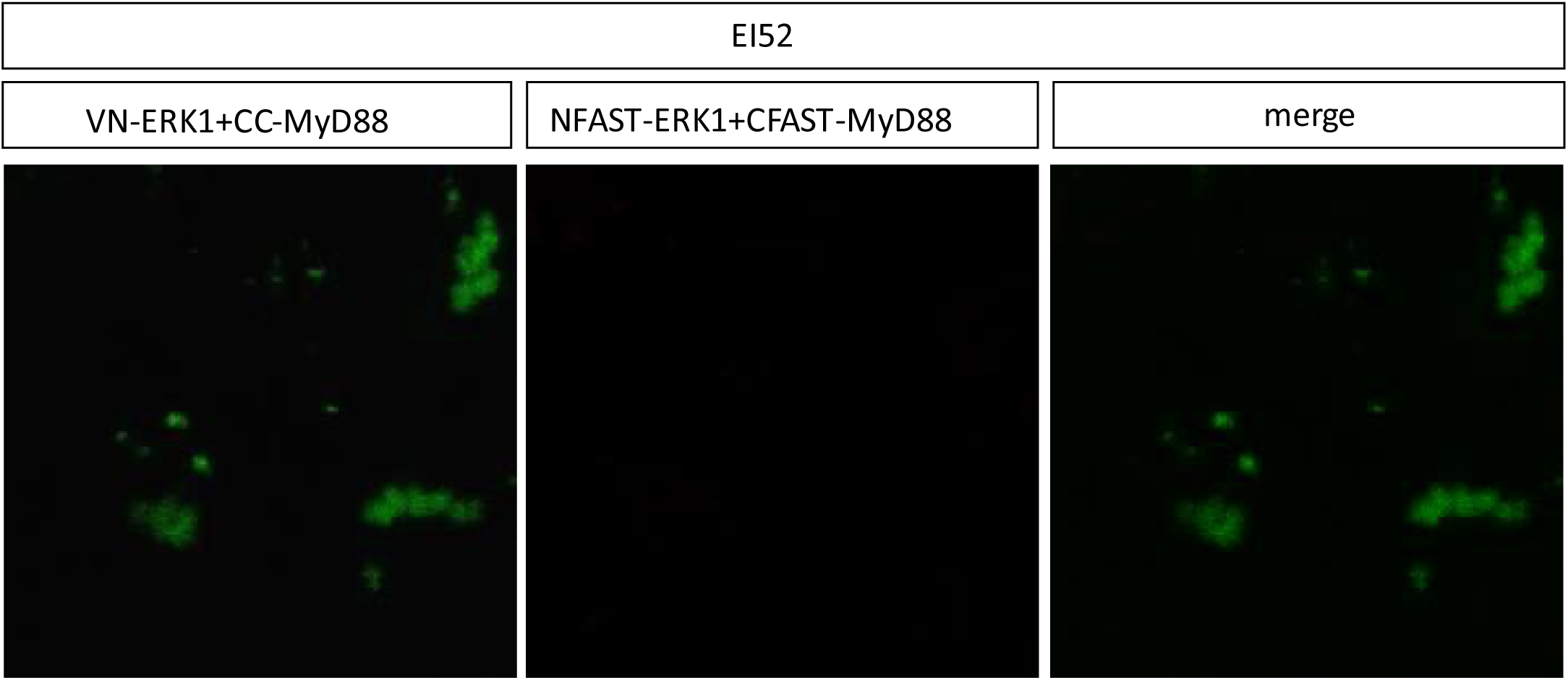
BiFC and splitFAST2 of ERK1/MyD88 interaction in the presence of DMSO+EI52 from 0 to 10 minutes after the addition of the HBR ligand. DMSO+EI52 that was added 12h post-transfection and cells were cultured for additional 6h in the presence of DMSO+EI52 before live imaging.

